# Pharmacogenomic synthetic lethal screens reveal hidden vulnerabilities and new therapeutic approaches for treatment of NF1-associated tumors

**DOI:** 10.1101/2024.03.25.585959

**Authors:** Kyle B. Williams, Alex T. Larsson, Bryant J. Keller, Katherine E. Chaney, Rory L. Williams, Minu M. Bhunia, Garrett M. Draper, Tyler A. Jubenville, Wendy A Hudson, Christopher L. Moertel, Nancy Ratner, David A. Largaespada

## Abstract

Neurofibromatosis Type 1 (NF1) is a common cancer predisposition syndrome, caused by heterozygous loss of function mutations in the tumor suppressor gene *NF1*. Individuals with NF1 develop benign tumors of the peripheral nervous system (neurofibromas), originating from the Schwann cell linage after somatic loss of the wild type *NF1* allele, some of which progress further to malignant peripheral nerve sheath tumors (MPNST). There is only one FDA approved targeted therapy for symptomatic plexiform neurofibromas and none approved for MPNST. The genetic basis of NF1 syndrome makes associated tumors ideal for using synthetic drug sensitivity approaches to uncover therapeutic vulnerabilities. We developed a drug discovery pipeline to identify therapeutics for NF1-related tumors using isogeneic pairs of *NF1-*proficient and deficient immortalized human Schwann cells. We utilized these in a large-scale high throughput screen (HTS) for drugs that preferentially kill *NF1*-deficient cells, through which we identified 23 compounds capable of killing *NF1-*deficient Schwann cells with selectivity. Multiple hits from this screen clustered into classes defined by method of action. Four clinically interesting drugs from these classes were tested *in vivo* using both a genetically engineered mouse model of high-grade peripheral nerve sheath tumors and human MPNST xenografts. All drugs tested showed single agent efficacy in these models as well as significant synergy when used in combination with the MEK inhibitor Selumetinib. This HTS platform yielded novel therapeutically relevant compounds for the treatment of NF1-associated tumors and can serve as a tool to rapidly evaluate new compounds and combinations in the future.

## Introduction

Neurofibromatosis Type 1 (NF1) is a cancer predisposition syndrome and one of the most prevalent autosomal dominant genetic disorders, occurring in one in every 2,500 individuals, with over 100,000 affected people in the United States alone^1,2^. Individuals with NF1 exhibit a wide variety of manifestations including tumors, café-au-lait macules, freckling, vasculopathy, skeletal dysplasia, and learning disabilities. Importantly, individuals with NF1 are at an increased for cancer in general, with a recent study showing that the lifetime risk of all cancers for NF1 patients at ∼60%^3^. The wide range of disease manifestations and the diverse range of severity of NF1 presentation among individuals make NF1 a complex whole-body disorder that is difficult to study.

NF1 is characterized by mutations in the tumor suppressor gene *NF1*, causing patients to develop benign Schwann cell tumors of the peripheral nervous system called neurofibromas^1,4^. These can progress to malignant peripheral nerve sheath tumors (MPNSTs), a deadly soft tissue sarcoma^5^. In fact, MPNSTs are the most common cause of disease associated death in NF1^6,7^. Current treatment options for MPNSTs show limited efficacy, with the only curative treatment being complete surgical resection^8,9^. MPNSTs often show local recurrence, and frequently metastasize^10^. While traditional cytotoxic chemotherapeutics are used to treat MPNST, they do not reduce mortality and offer only modest delays in disease progression or death^11,12^. The genetic basis of NF1 syndrome makes it an ideal candidate for using synthetic lethal genetic and therapeutic approaches to uncover unique variabilities specific to *NF1*-deficient cells. Identification of new molecular targets for therapeutics effective against both benign tumors and MPNSTs will be critical for improved patient outcomes and quality of life.

Understanding cellular adaptations and new molecular vulnerabilities that accompany loss of both copies of the *NF1* gene in relevant cell types is predicted to reveal novel therapeutic opportunities. Therefore, we chose to utilize synthetic drug lethality to uncover novel therapeutic vulnerabilities found exclusively in *NF1-*deficient human Schwann lineage cells (Figure 1). One copy of the *NF1* gene harbors a loss of function mutation in every cell of an NF1 patient; both neurofibromas and MPNSTs result from loss of heterozygosity (LOH) for the second copy of *NF1*^13^. This presents a scenario analogous to *BRCA1/2* mutated cancers, such as breast and ovarian carcinomas^14,15^, in which synthetic lethality has been established as a clinically relevant paradigm that revealed PARP inhibitors exploit the DNA repair pathway vulnerability intrinsic to those cells^16^ (Figure 1A).

**Figure 1.**
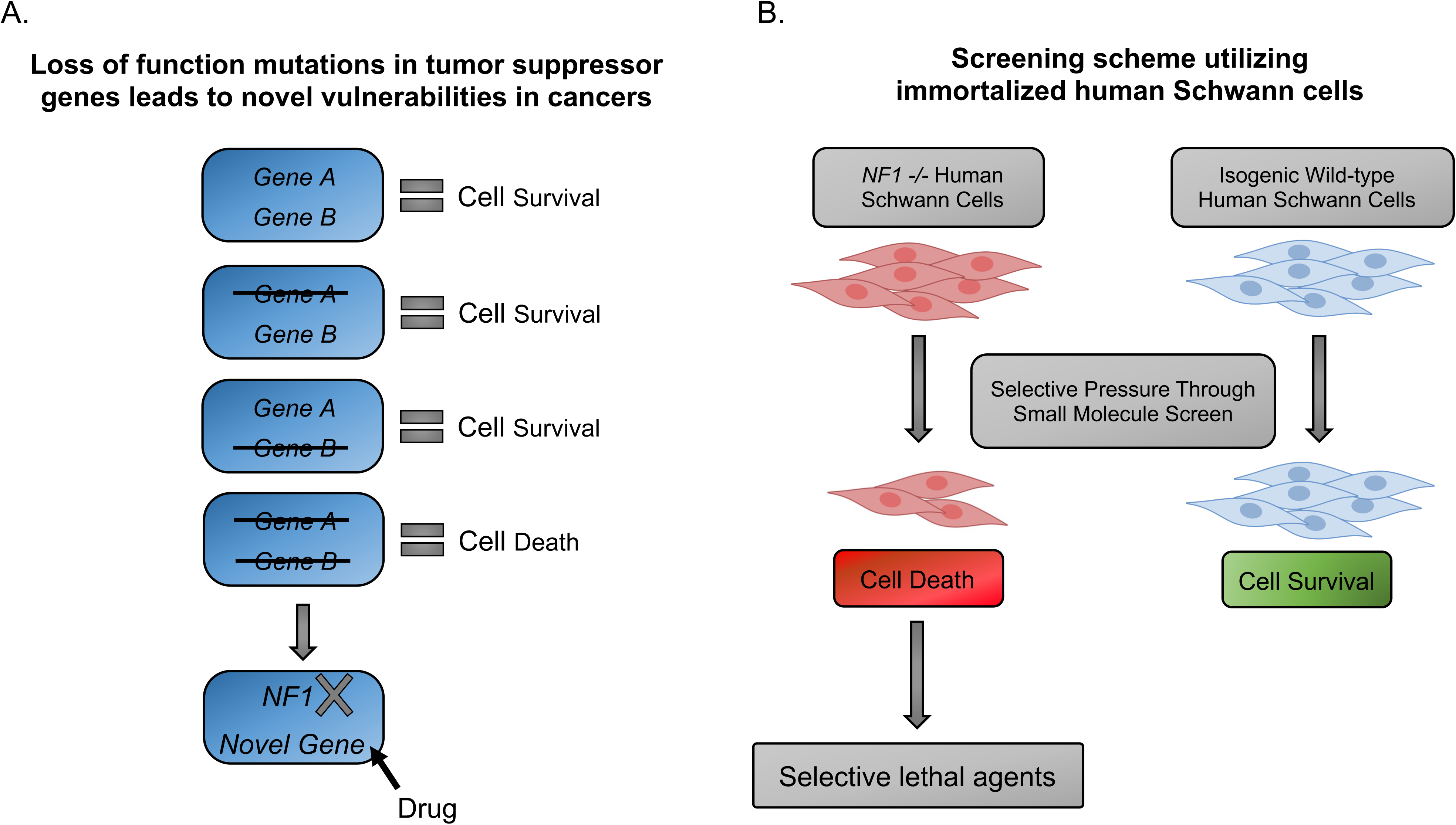
Targeting cell intrinsic vulnerabilities convened by loss of tumor suppressor gene function. A. Synthetic lethality is the genetic incompatibility of the loss of two or more gene products within the same cell. In cells already lacking functional tumor suppressors, such as *NF1*, it is possible to target the function of a second protein to induce synthetic drug lethality. B. Identifying selective lethal therapeutic agents effective against *NF1*-deficient human Schwann cells via selective pressure from a large-scale small molecule screen.

Several drug screening projects have been previously undertaken for NF1, with some success. Those projects relied on using mouse embryonic fibroblasts mutated for the *NF1* gene, or yeast cells, or MPNST cell lines ^17–20^. These approaches each have limitations. For instance, if cell type and context are important for drug sensitives and synthetic lethal interactions, Schwann cell lineage cells may be critical to identify relevant targets ^21–24^. MPNST cell lines are of the relevant lineage, but there are few MPNST cell lines in use by the research community so the available lines may not reflect the genetic heterogeneity of patients’ tumors. Also, each MPNST cell line likely has many passenger mutations, copy number variants, and varying degrees of aneuploidy, even for the “same” cell line depending on how long it has been in culture, as noted in other cancer models ^25^. Moreover, cancer cell lines are fast growing and are generally susceptible to death by traditional chemotherapeutics, drugs that have limited effectiveness against MPNST^12^. Cell type and context can also be important for drug sensitives and synthetic lethal interactions^21,22,24^, these are lost if not working with the cell type of origin for NF1-associated tumors.

Given that both plexiform neurofibromas and MPNSTs arise within the Schwann cell lineage, we developed a drug discovery pipeline to identify targeted therapeutics for NF1-related neoplasia, including MPNSTs. In our study, we used CRISPR-Cas9 genome editing techniques to introduce specific *NF1* mutations into immortalized human Schwann cells, and then screened a large and diverse library of drugs to identify those that can selectively kill the *NF1-*deficient cells (Figure 1B). This approach allowed us to identify drugs that target the specific genetic mutations found in NF1 associated tumors, making them effective against these tumors while minimizing the toxicity to normal cells ^26^.

Using high throughput drug screening (HTS), we identified several compounds and drug classes that are selective against *NF1-*deficient human Schwann cells. Of these, two were of particular clinical interest, as they are either FDA approved for other indications or are in late-stage clinical development with some Phase III data available. Digoxin, a long-used cardiac glycoside^27,28^ was particularity potent against *NF1-*deficint human Schwann cells and MPNST cell lines. Moreover, digoxin synergized with the MEK inhibitor Selumetinib, which is FDA approved for treatment of symptomatic NF1-associated plexiform neurofibromas (PN)^29,30^. The microtubule destabilizing agent rigosertib^31^ showed similar specificity and efficacy, both alone and in combination with Selumetinib, in our study. Both drug combinations exhibited efficacy in a genetically engineered mouse model of high grade peripheral nerve sheath tumors^32^ and in xenograft models of MPNST^33^, and showed the ability to both dramatically shrink some established tumors and give complete responses in other animals. These results identify novel combination drug therapies that target NF1-related neoplasia, including MPNST, using drugs which can be rapidly tested for efficacy in NF1 patients.

## Materials and Methods

### Tissue culture

Immortalized human Schwann cell lines were maintained and passaged at 37°C with 5% CO_2_ in DMEM high glucose media supplemented with 10% FBS and penicillin/streptomycin. If antibiotic selection was needed, puromycin was used at 500 µg/ml as indicated below. S462-TY cells were cultured in DMEM supplemented with 10% FBS and 1% penicillin-streptomycin at 37°C and 5% CO_2_. Mycoplasma detection was routinely performed using the MycoAlert detection kit (Lonza) throughout this study. Cell line authentication was performed at the University of Arizona Genetics Core against established reference lines.

### Generation of *NF1-*deficient cell lines

We received immortalized human Schwann cell (SC) lines (iHSCs) from Dr. Margaret Wallace (University of Florida, Gainesville). These are wild-type human SCs immortalized using human reverse transcriptase component of telomerase (hTERT) and murine cyclin dependent kinase (Cdk4) transgenesis^11^. All subsequent mutations were engineered into this cell line. U6-gRNA vectors were produced as previously described^34^ and introduced into iHSCs that are proficient for *NF1*. U6-gRNA vector was engineered with unique sequences targeting a site in *NF1* to cause insertion-deletion (indel) mutations in a critical exon to introduce frameshift mutations. The target sequences can be found in Figure 2A.

**Figure 2.**
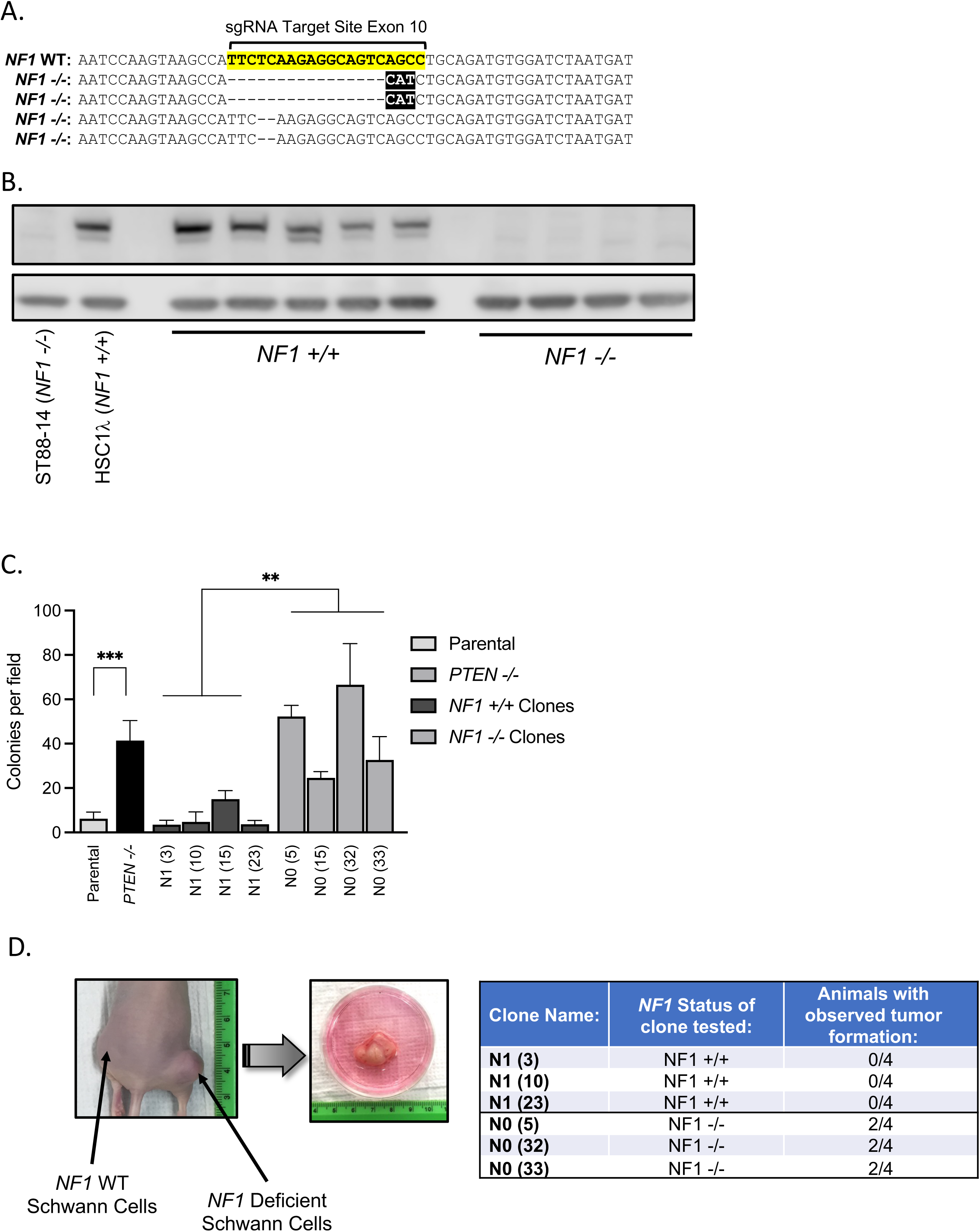
Characterization of *NF1-*deficient immortalized human Schwann cells. A. Exon 10 of the *NF1* gene was targeted for mutagenesis using CRISPR-Cas9. The sgRNA target site in the wild type *NF1* sequence is highlighted in yellow. Independent clones were recovered harboring the indicated biallelic indels at this locus which result in frameshifts and early stop codons. B. Immunoblot showing the immortalized human Schwann cell lines identified as having biallelic loss of function mutations in *NF1* do not make neurofibromin protein. *NF1 +/+* samples come from isogenic sister clones to the *NF1-*deficient cells which when sequenced did not have mutations at the guide target site in *NF1*. HSC1l is the parental cell line and serves as a positive control. ST88-14 is an established *NF1-*deficient MPNST cell line. C. *NF1-*deficient human Schwann cells form significantly more colonies in soft agar under low serum conditions compared to isogenic matched *NF1-*proficient lines and are poised for transformation. Parental HSC1l cells and those in which the tumor suppressor *PTEN* has been knocked out serve as controls. ** *p*<0.01, *** *p*<0.001. D. *NF1-*deficent human Schwann cells *can* form xenograft tumors in the flanks of immunodeficient athymic nude mice. 1×10^6^ cells of 3 *NF1-*proficient and 3 *NF1*-deficient clones were each implanted in the flanks of 4 mice and monitored for tumor development. *NF1*-deficient clones showed the ability to form tumors while tumor formation was not observed for the *NF1*-proficient clones.

All vectors were introduced via electroporation and performed using the NEON electroporation system (Invitrogen) using 100µL electroporation tips according to the manufacture’s protocol. One million cells were electroporated (1 pulse of 1400V for 30ms) with 2 µg Flag-hCas9, 2ug each U6-gRNA vector (gifts from Branden Moriarity), and 100ng pmaxGFP plasmid (Amaxa) to assess transfection efficiency^34^. To enrich for modified cells, co-transposition was performed using 500 ng of the *piggyBac* (*PB*) transposon PB-CAGG-Luciferase-IRES-GFP-PGK-Puro^R^, and 500 ng of CMV-*PB7 transposase*^34^. Cells were provided fresh growth medium one day after electroporation, and two days later were selected with puromycin (500 µg/mL; Fisher) for one week. Selective medium was changed every 2-3 days for one week.

Limiting dilutions were then plated from the pooled cell lines, which are enriched for cells edited at the sites of interest, to obtain single-cell clones. These clones were then expanded for future work and validation. Validation of the mutations was done by obtaining genomic DNA from the derived clones, amplifying DNA around the gRNA target/cut sites, and subjecting to Sanger sequencing. Amplicons were generated in 50 µL reactions with GoTaq Green Master Mix (Promega) using the following PCR cycle: initial denaturation at 95°C for 5 min; 30 × (95°C for 30sec, 59°C for 35sec, 72°C for 2min); final extension at 72°C for 5 min. Forward (AACAGCTTGTTTGGGAAGGA) and reverse (CATTGGTGATGATTCGATGG) primers amplified a 1323bp region flanking *NF1* exon 10. PCR products were purified and sequenced using a third primer binding in *NF1* intron 9-10 (TGGCAGCTGGATTTTACTGC). Clones targeted for editing of exon 10 in *NF1* were recovered with a variety of deletions/insertions around the target site, verified through directed Sanger sequencing. A summary of recovered clone sequences is shown in Figure 2A. Western blot analysis was also done on a subset of clones of interest to validate functional knockout of neurofibromin expression (Figure 2B).

### *In vivo* tumor forming ability of iHSCs

*NF1-*proficient and *NF1-*deficient engineered clones were injected into the flanks of 6-to 7-week-old athymic nude mice (1.0×10^6^ cells) in 50% Matrigel (BD Biosciences). Mice were monitored for tumor formation for 6 months post injection or until a tumor volume of 2000mm^3^ was reached.

### Western blotting

Immunoblotting was done on whole cell lysates collected from various cell lines. After collection, cells were washed with 1x PBS, collected, pelleted and flash frozen. Pellets were then thawed and lysed using 1x RIPA buffer (Sigma) supplemented with phosphatase inhibitor cocktails 2 & 3 (Sigma). Whole cell lysates were vortexed at 2,000 RPM for 15 minutes before being sonicated with 30-1 second pulses. Then, 10-30ug of whole cell lysate was loaded into each well of 4-12% Bis-Tris NuPage polyacrylamide gels (Invitrogen). Proteins were transferred overnight onto PVDF membranes. Each membrane was blocked with 5% BSA in TBST for 1 hour and incubated with 1:1000 primary antibody overnight at 4°C. After rinsing away primary antibodies, membranes were incubated for 1 hour with 1:5000 secondary antibody. WesternBright Quantum HRP substrate (Advansta) was added to membranes before being imaged on an Odyssey Fc imager (LI-COR Biosciences). Primary and secondary antibodies used were from Cell Signaling Technology: Neurofibromin 1 (D7R7D) Rabbit mAb 14623 anti-rabbit IgG and HRP-linked secondary 7074 (CST). β-actin was probed as a loading control using β-Actin (13E5) Rabbit mAb #4970 (Cell Signaling Technologies) at 1:5,000.

### Small molecules and drug libraries

All drug screening libraries used in this study were obtained from the University of Minnesota Institute for Therapeutics Design and Development collection. Validation of hits from the primary drug screening effort and all *in vivo* work was done with repurchased compounds from SelleckChem.

### *In vitro* combination drug testing and analysis

Cells were seeded into 384-well plates at 2,000 cells/well using a Biomek 2000. After 24hr of growth in medium supplemented with 10% FBS and penicillin/streptomycin, drug was added in triplicate or quadruplicate wells per dose in a 8-point or 12-point dose-response manner (depending on experiment, as indicated) using the acoustic Echo 550 liquid dispenser (Labcyte). Drug combination testing was done in a similar manner and plated in a constant ratio as indicated. After incubation with drug(s) or vehicle for 48 hrs, cells were incubated with alamarBlue (Thermo Fisher) reagent and fluorescence read on a CLARIOstar microplate reader (BMG LABTECH). Cell viability was calculated by fluorescence of experimental wells in percent of unexposed control wells with blank values subtracted. Data were analyzed in Prism software (Graphpad Prism) and dose-response curves generated using nonlinear regression log(inhibitor) vs. response-variable slope model. Each point represents mean ± SD.

### Combination index value determination

Drug interaction values were determined using the median-effect principle of Chou-Talalay^35,36^. Combination index (CI) values were calculated using CalcuSyn software (Biosoft) as previously described^37^. CI<1, =1, and >1 indicate synergism, additive effect, and antagonism, respectively.

### Transcriptional response to therapeutics

The S462TY MPNST cell line was grown under standard tissue culture conditions. Cells were exposed to vehicle (DMSO), digoxin, rigosertib, selumetinib, digoxin + selumetinib, or rigosertib + selumetinib for 2 hours or 24 hours. For drug treatments, all compounds were used at the following concentrations: selumetinib [125 μM], rigosertib [0.5 μM], digoxin [0.0308 μM], vorinostat [2.5 μM], selumetinib [15.625μM] plus vorinostat [1.56 μM], selumetinib [4 μM] plus digoxin [0.0308 μM], and selumetinib [2 μM] plus rigosertib [0.1 μM]. At the indicated time point, cells were harvested, washed with cold PBS, pelleted via centrifugation, and the pellets flash frozen in LiN_2_. Cell pellets were stored at –80°C until RNA purification.

### RNA Purification and Sequencing

RNA was harvested from cells using Qiagen RNEasy mini purification kits and subjected to TURBO DNAse to remove any residual genomic DNA contamination. RNA quality was assayed via Agilent Bioanalyzer and sequenced on an Illumina NovaSeq 6000 next generation sequencer.

### RNAseq Analysis

RNAseq data was analyzed as follows: Quality trimming via Trimmomatic (v0.33)^38^, alignment via HISAT2 (v2.1.0)^39^ to the GRCh38 Ensembl assembly^40^, and gene count quantification via StringTie(v1.3.4d)^41^. Differential expression analysis was performed using R Statistical Software(v4.3.1)^42^, tidyverse(v2.0.0)^43^ DESeq2(v1.42.0)^44^ Pathway analysis was performed using Gene Analytics(v)^45^. The data discussed in this publication have been deposited in NCBI’s Gene Expression Omnibus^46^ and are accessible through GEO Series accession number GSE262030 (https://www.ncbi.nlm.nih.gov/geo/query/acc.cgi?acc=GSExxx).

### Flow cytometry

Cell cycle stage and apoptosis was determined by staining and flow cytometry. S462-TY cells were treated with drug compounds or DMSO control for 0, 24 and 48 hours before harvest. For cell cycle stage determination, cell supernatant and adhered cells were collected and stained for flow cytometry using a Propidium Iodide kit (Abcam ab139418) using manufacturer’s protocol. Briefly, after collection, cells were washed with 1x PBS then fixed using ice cold 66% ethanol. Cells were then washed again with 1x PBS and stained with 1x Propidium Iodide and RNase Staining Solution before being run through the CytoFLEX flow cytometer (Beckman Coulter). Cell cycle stage was determined with the Cell Cycle tool in FlowJo V10.7.1 Software (BD Life Sciences) using the Watson (Pragmatic) model.

### *In vivo* models and drug testing

For the MPNST cell line xenograft model, S462-TY cells (1.0×10^6^) in 50% Matrigel (BD Biosciences) were injected subcutaneously into the flank of 6 to 7-week-old NOD-Rag1null IL2rgnull, NOD rag gamma, NOD-RG (NRG) mice. For the patient derived xenograft (PDX) MPNST model, tumor tissue was implanted subcutaneously directly into NOD.Cg-Prkdc^scid^ Il2rg^tm1Wjl^/SzJ (NSG) mice^33^. Administration of drugs was started when tumors reached 150–200 mm^3^. All treatments were administered via intraperitoneal (IP) injection at the following doses: digoxin 2mg/kg, rigosertib 100mg/kg, selumetinib, 10mg/kg.

The genetically engineered mouse model of high-grade peripheral nerve sheath tumors (PNSTs) was generated as described previously^32^. *Dhh::Cre, Nf1^fl/fl^, Pten^fl/fl^*experimental class animals were randomized to a treatment or vehicle group at 1 week of age. All treatments were administered via IP injection.

## Results

### Cell line characterization

Exon 10 of the *NF1* gene was targeted for CRISPR/Cas9 mediated mutagenesis in the immortalized human Schwann cell line HSC1l. Clones were recovered harboring biallelic loss of function insertion/deletion (indel) mutations with the sequence indicated in Figure 2A. A number of sister clones that also went through the mutagenesis process were also recovered in which no identifiable mutations in *NF1* were found. These *NF1-*proficient clones produce full length neurofibromin, while the *NF1*-deficient clones lack detectible neurofibromin expression (Figure 2B). *NF1*-deficient clones also show increased basal RAS-GTP levels, as expected, when compared to the isogenic matched *NF1-*proficient sister clones (Supplementary Figure 1).

*NF1-*deficient human Schwann cells exhibit an increase in transformed phenotypes. Under low serum conditions the *NF1-*deficient cells showed a significantly greater capacity for anchorage independent growth. Parental HSC1l and *NF1-*proficient cells form limited numbers of colonies when grown in low-serum soft agar. All *NF1*-deficient lines tested exhibited robust colony formation in these conditions, similar to the phenotype of an HSC1l cell line in which the tumor suppressor *PTEN* has been knocked out (Figure 2C).

The *NF1-*deficient cell lines are poised for transformation. Three *NF1*-proficient and three *NF1-* deficient human Schwann cell lines were tested for the ability to grow as xenografts in the flanks of immunodeficient athymic nude mice. Each line was injected into the flanks of 4 mice and monitored for signs of tumor formation for 16 weeks. Tumor formation was never observed for the *NF1-*proficient cell lines. In contrast, all *NF1-*deficient lines showed the ability to form tumors upon xenograft, with 50% penetrance (Figure 2D). When the same lines were tested in less stringent NRG mouse strain the *NF1-*deficient lines developed tumors with 100% penetrance (Supplemental Table 1).

### In vitro high throughput drug screening

A panel of 11,085 small molecules was screened in an HTS pipeline. Candidates for testing were chosen from existing drug/small molecule libraries enriched for FDA approved drugs, drugs with some level of clinical development, and drug like compounds, to aid in translation of promising findings to the clinic. Table 1 lists the libraries screened.

**Table 1.** Libraries Screened.

| **Library:** | **Number of**<br>**Compounds:** | **Reference/Source:** |
| --- | --- | --- |
| **General:** |  |  |
| NIH Clinical Collection | 446 | www.nih.gov |
| Prestwick | 1120 | www.prestwickchemical.com |
| Tocris Screening | 1120 | www.tocris.com |
| Microsource Spectrum | 2000 | www.msdiscovery.com |
| Johns Hopkins | 1514 | drugdiscovery.jhu.edu |
| NCI Set | 2396 | www.nci.org |
| Sigma LOPAC | 1280 | www.sigmaaldrich.com |
| CTF/SYNODOS | 12 | *This work* |
| **Kinase Specific:** |  |  |
| Tocris Kinase | 80 | www.tocris.com |
| GSK Kinase Inhibitor Set 1 | 367 | itdd.umn.edu |
| GSK Kinase Inhibitor Set 2 | 473 | itdd.umn.edu |
| SelleckChem Kinase | 277 | www.selleckchem.com |
| **Total:** | **11085** |  |

All 11,085 compounds were tested in triplicate against the *NF1−/−* cell line N0 (5) at a single concentration of 10μM. ∼650 compounds were identified as being active against this cell line, by inhibiting viability >= 50% after 48 hrs. of growth (Figure 3A). These primary screening hits were then tested across an 8-or 12-point dose response range against an *NF1-*proficient, N1 (10), and an *NF1-*deficient cell N0 line (5). This secondary screening step identified 27 drugs that were selectively lethal to the *NF1*-deficient cell line. Many of these drugs grouped together in classes with similar method of action, including topoisomerase inhibitors, endoplasmic reticulum (ER) stress/unfolded protein response (UPR) stimulation, NF-κB pathway modulation, cytoskeletal perturbation, MEK inhibition, and intracellular calcium modulators (Table 2).

**Figure 3.**
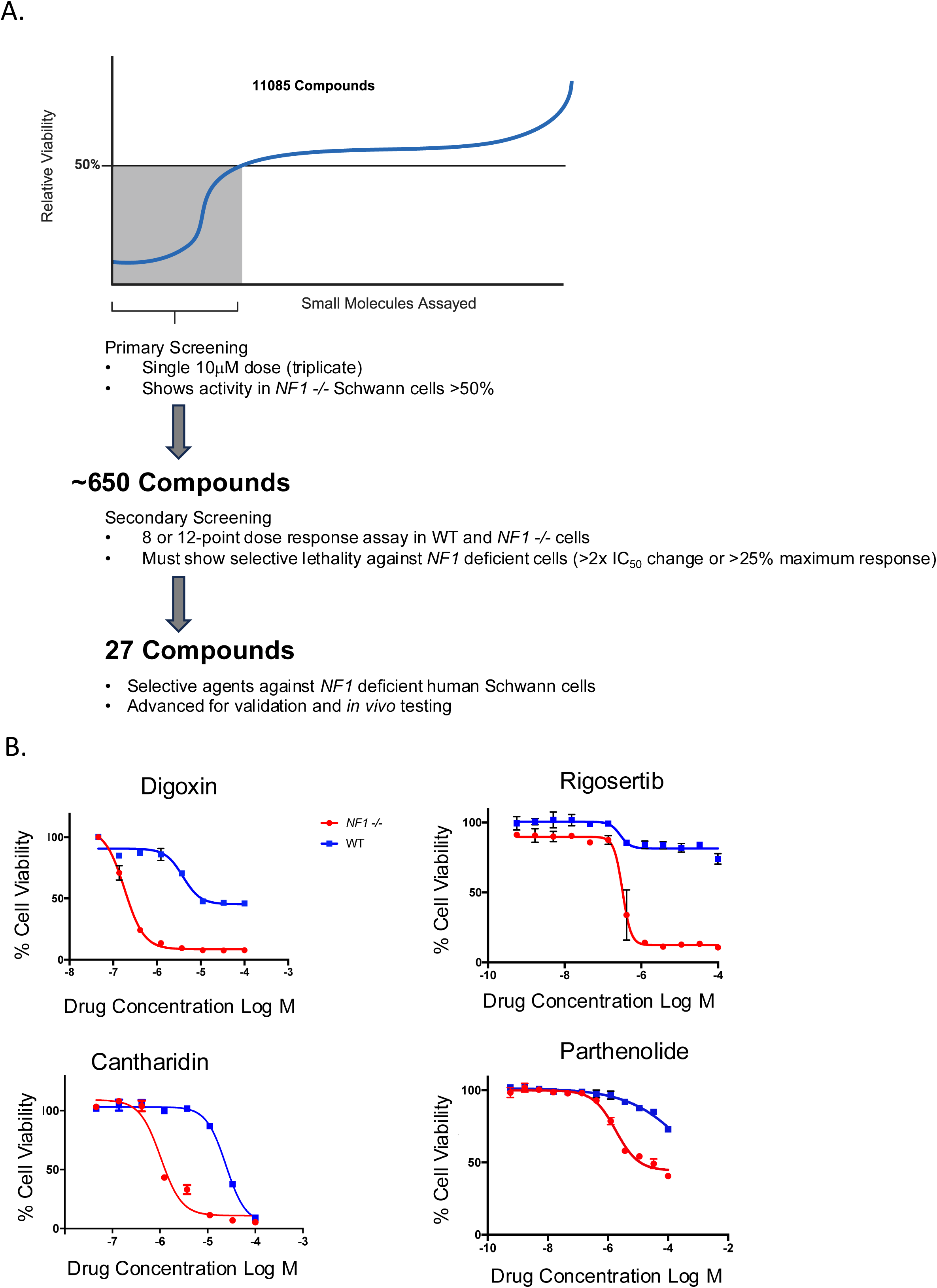
High throughput screening (HTS) of small molecule libraries identifies compounds selectively lethal to *NF1*-deficient human Schwann cells. A. Over 11,000 compounds were used for primary screening against *NF1*-deficient clones to identify active molecules. This was done in triplicate at a single dose of 10μM. Compounds showing a reduction of viability of ≥50% were considered hits. ∼650 hits from primary screening were then tested in dose response assays against both *NF1*-profieient and *NF1*-deficient isogenic lines to identify selective lethal compounds. This identified 27 compounds found to be selective against *NF1*-deficient Schwann cells that were advanced for validation and *in vivo* testing. B. Dose response curves for 4 clinically interesting compounds identified in this screen. Each drug was tested against *NF1*-deficient and proficient isogenic matched clones across a wide range of concentrations. Large shifts in IC_50_ and overall efficacy are observed among these candidates.

**Table 2.**
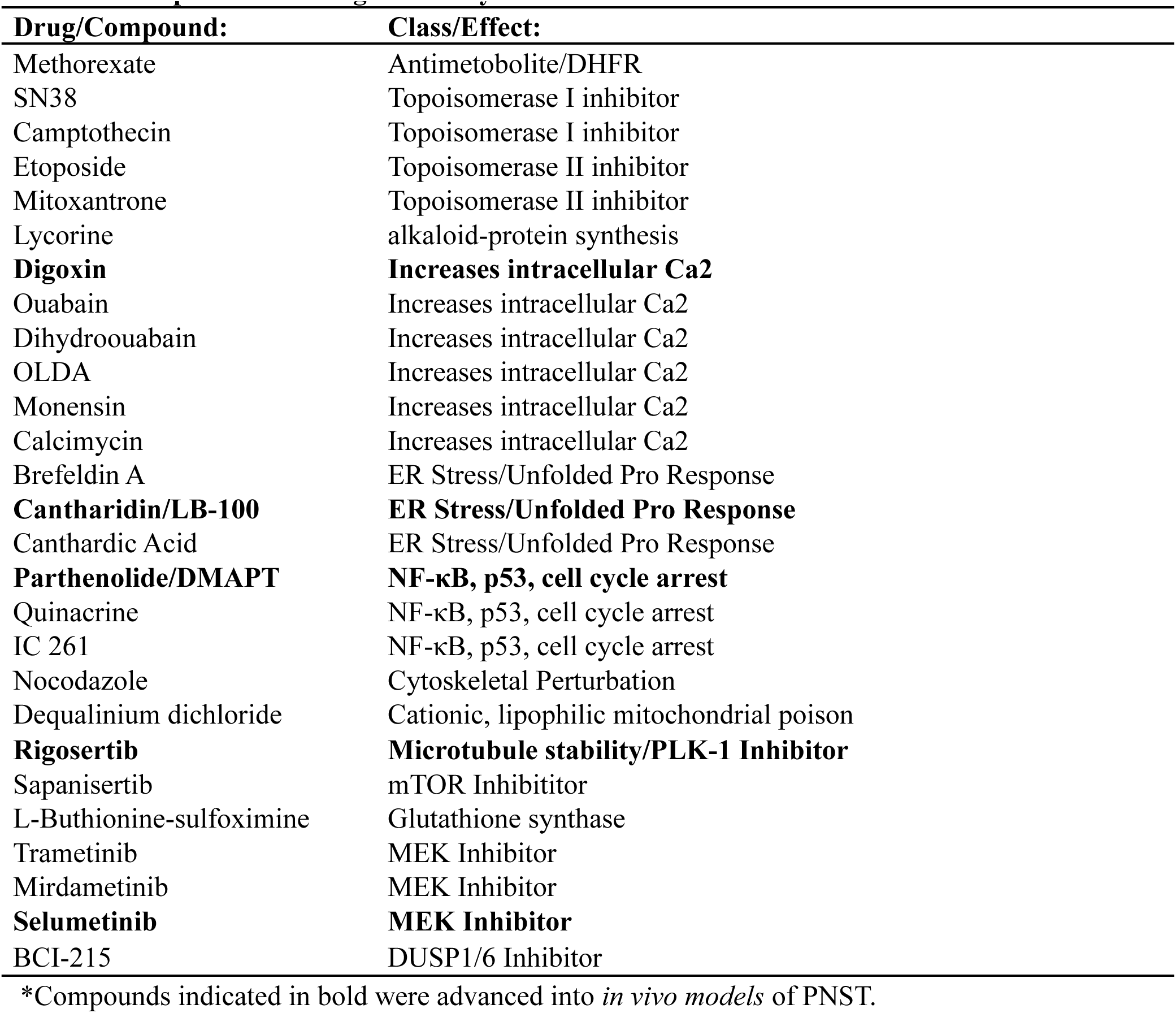
Compounds showing selectivity for inhibition of *NF1-*defcient Schwann cells.

Drug selectivity for *NF1 −/−* cells was defined as lower IC_50_ concentrations and/or increased overall efficacy (reduced cell viability), as compared to effects in wild type cells. The small molecules digoxin, rigosertib, cantharidin, and parthenolide represent hits from four of the key target classes identified in the primary screen that show selectivity toward the *NF1-*deficient cells (Figure 3B). Many compounds capable of increasing intracellular calcium were also selective against *NF1* −/− cells (Table 2). For example, the cardiac glycoside digoxin, which is clinically used to treat congestive heart failure, exhibited both a marked shift in IC_50_ (∼1 log) and increased potency (∼40%) toward *NF1* −/− cells.

Two of the highly selective and potent drugs (digoxin and rigosertib) from the HTS effort had clinical translatability. To verify applicability for the treatment of these malignancies, we tested these drugs against a panel of MPNST cell lines. As when digoxin was used against *NF1-*deficient immortalized human Schwann cells, a shift of over 1 log was seen in the IC_50_ when comparing the MPNST cell line S462-TY to *NF1-*proficient immortalized human Schwann cells, with a near doubling of potency (Fig 4A). The microtubule destabilizing agent Rigosertib showed a 50% increase in potency against S462-TY cells. Rigosertib showed an IC_50_ in the nM range for S462-TY cells, but was not effective against *NF1-*proficient immortalized human Schwann cells (Fig 4A).

**Figure 4.**
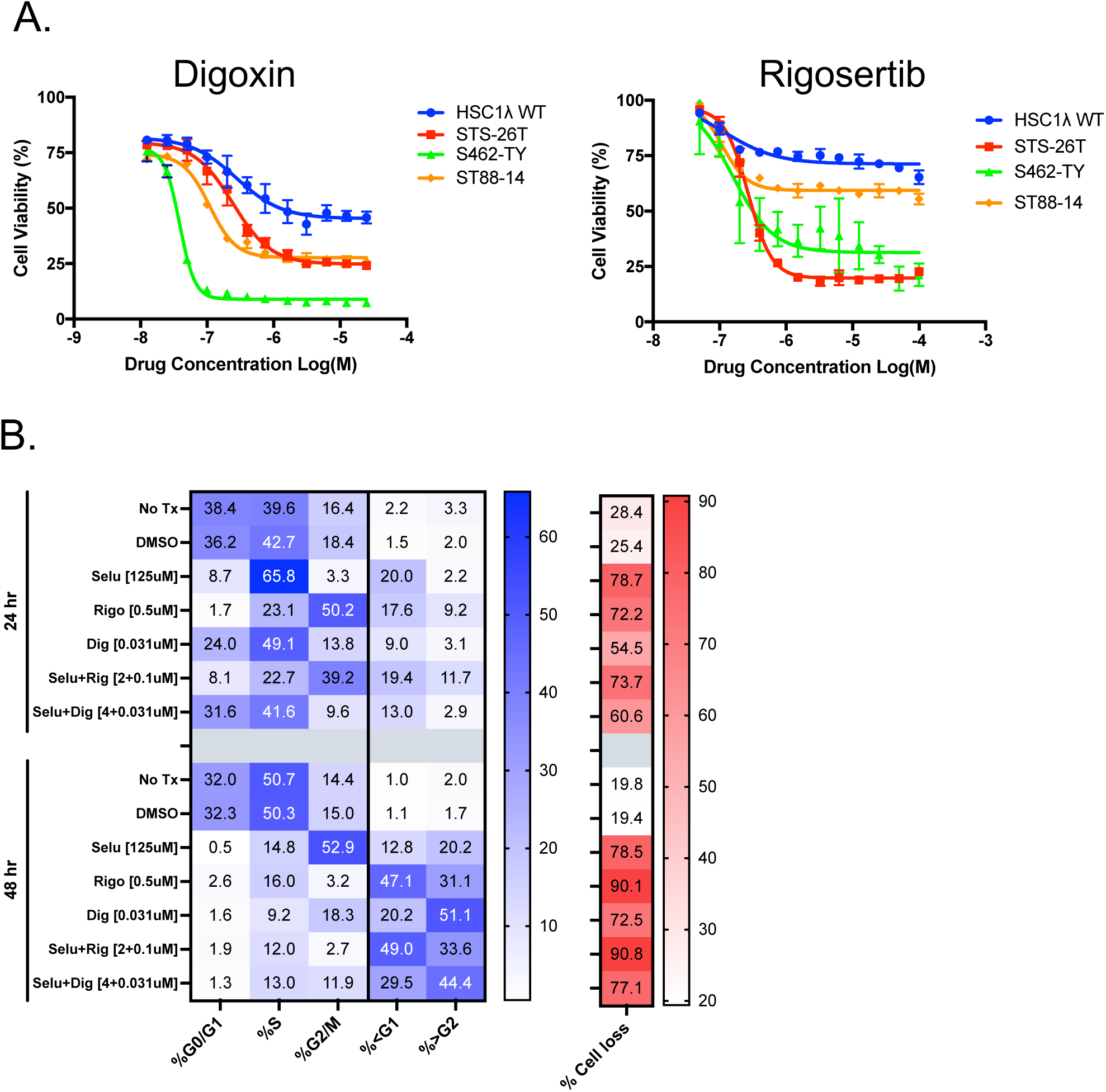
Selected drugs shown to be effective and selective against *NF1 −/−* show effectiveness against human MPNST cell lines and synergize with MEK inhibition. A. 12-point dose response curves of digoxin (left) and rigosertib (right) against several human MPNST cell lines. *NF1*-proficient HSC1l serve as a control. B. Cell cycle analysis of S426-TY MPNST cell line treated with selumetinib, rigosertib, digoxin as single agents and in combination at 24 and 48 hrs.

### Synergy with MEK inhibition

Rigosertib and digoxin demonstrated significant selective killing of *NF1*-deficient human Schwann cells and MPNST, as well as significant efficacy (Figure 4A). Rigosertib is currently in multiple late-stage clinical trials for cancer therapy^47,48^ and digoxin has been FDA approved for other indications for many years^49,50^. These clinically interesting drugs were tested in combination with the MEK inhibitor Selumetinib as we hypothesized there would be synergistic killing effects with the combinations. Experiments to determine the Combination Index (CI) values of Selumetinib + rigosertib and Selumetinib + digoxin were conducted against a panel of cell lines including: *NF1-*proficient HSC1l, an *NF1-*deficient iHSC cell line, and two MPNST cell lines (S462-TY and ST88-14). The combination therapies were found to show synergy against *NF1−/−* cells, across a wide range of effective dose (ED) ranges (Table 3). Interestingly, the Selumetinib + digoxin combination showed positive CI values, suggesting an antagonistic interaction, and reduced killing ability against the *NF1-*proficient line, particularly at the ED^90^ level (CI 2.498). This suggests increased selectivity toward *NF1* mutant Schwann cells of the combination and perhaps predicts less toxic effects on normal tissues.

**Table 3.**
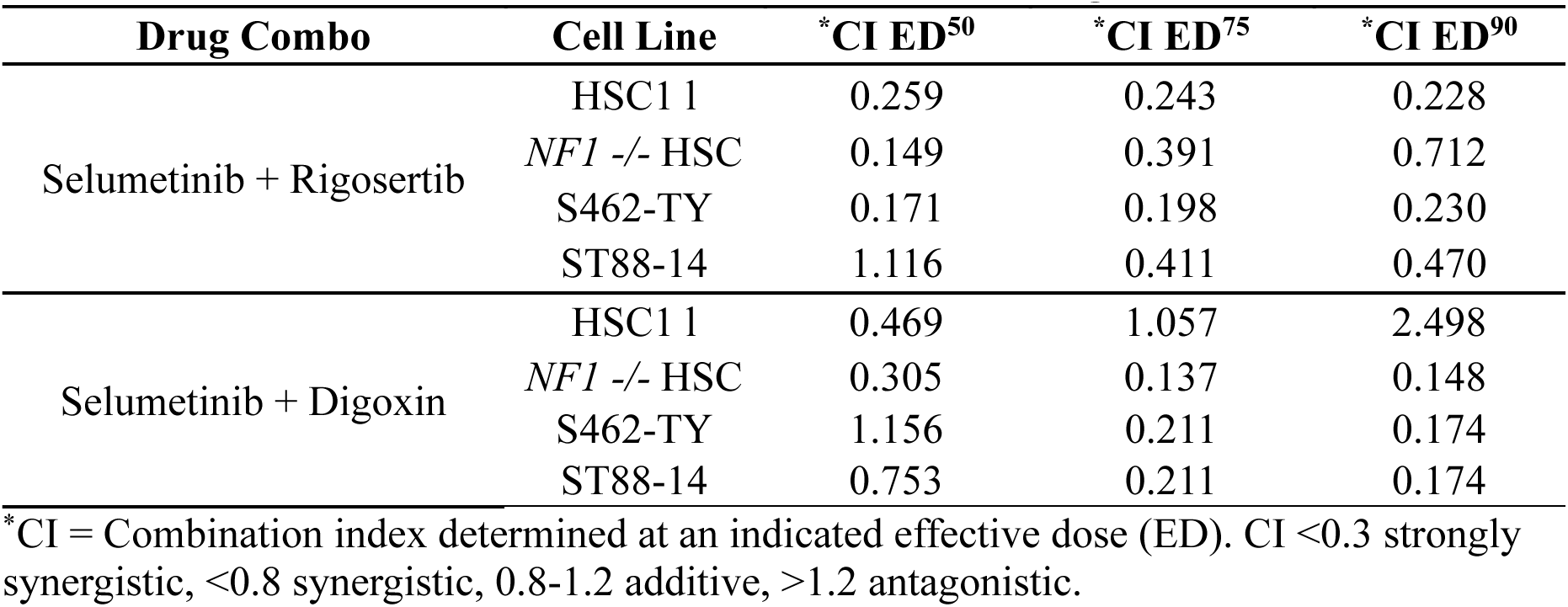
Combination index determination of candidate drugs with selumetinib.

Effects of rigosertib and digoxin on the cell cycle of S462-TY MPNST cells, alone and in combination with Selumetinib, were determined via flow cytometry (Figure 4 B, Supplementary Figure 2). S462-TY MPNST cells show limited sensitivity to MEK inhibition. At 24hrs, 125 μM Selumetinib alone results in ∼65% of the cells are showing an accumulation in S phase with only 8.7% in G0/G. Vehicle treated cells display 42.7% in S phase and 36.2% in G0/G1. At 24hrs 0.5μM rigosertib causes a G2/M arrest in the MPNST cell line (Figure 4B).

When selumetinib and rigosertib are combined, the predominate effect was a G2/M arrest with 39.2% of the cells residing in the G2/M population. However, this is achieved using much reduced drug versus single agents (2 μM vs 125 μM of selumetinib and 0.1 μM rigosertib vs 0.5 μM), to achieve similar cell cycle arrest and cell killing (Figure 4B). MEK inhibitors such as selumetinib can show significant side effects in patients, so opportunities to reduce the dose of a drug that could be a lifelong treatment is important to explore.

At 48 hrs of drug exposure the synergistic effects of the rigosertib + selumetinib combination were further increased. While 125 μM selumetinib alone produced the anticipated G2/M phase arrest, single agent treatment with rigosertib at 0.5 μM shows a dramatic shift of the population to <G1 (47.1%) and >G2 (31.1%) along with dramatic cell loss/killing. This suggests most of the remaining cells are having significant problems dividing or are in the process of dying. The combination treatment shows increased percentages of cells in these <G1 and >G2 populations, after treatment with the reduced drug concentrations.

Digoxin is highly potent against S462-TY. Only 31 nM was used in the 24 and 48 hr cell cycle analysis assays. At the 48 hr exposure, digoxin alone caused extensive cell loss and >50% of the remaining cells shifted to a >G2 population or a <G1 population. When used in combination with 4 μM selumetinib this same trend is seen, with an increase in the <G1 population to 29.5% of the cells.

### Novel cellular responses occur in human MPNST cells when exposed to combination therapy

To determine how the transcriptional profile of human MPNST cells changes when exposed to the drugs identified in this study, S462-TY MPNST cells were exposed to compounds for 24 hours after which RNA was harvested and subjected to RNA-sequencing. The gene expression fold change at 24 hours for each of the single agents (selumetinib, digoxin, and rigosertib) and the combinations (selumetinib + digoxin and selumetinib + rigosertib) versus the time matched vehicle control identified distinct transcriptional profiles of each treatment (Figure 5A). Each treatment condition resulted in many genes being upregulated or downregulated, and, importantly, new genes were regulated in response to the drug combination treatments, genes unchanged in the monotherapy responses. These emergent regulated genes represent the novel cellular response and can give insight into the observed drug synergies.

**Figure 5.**
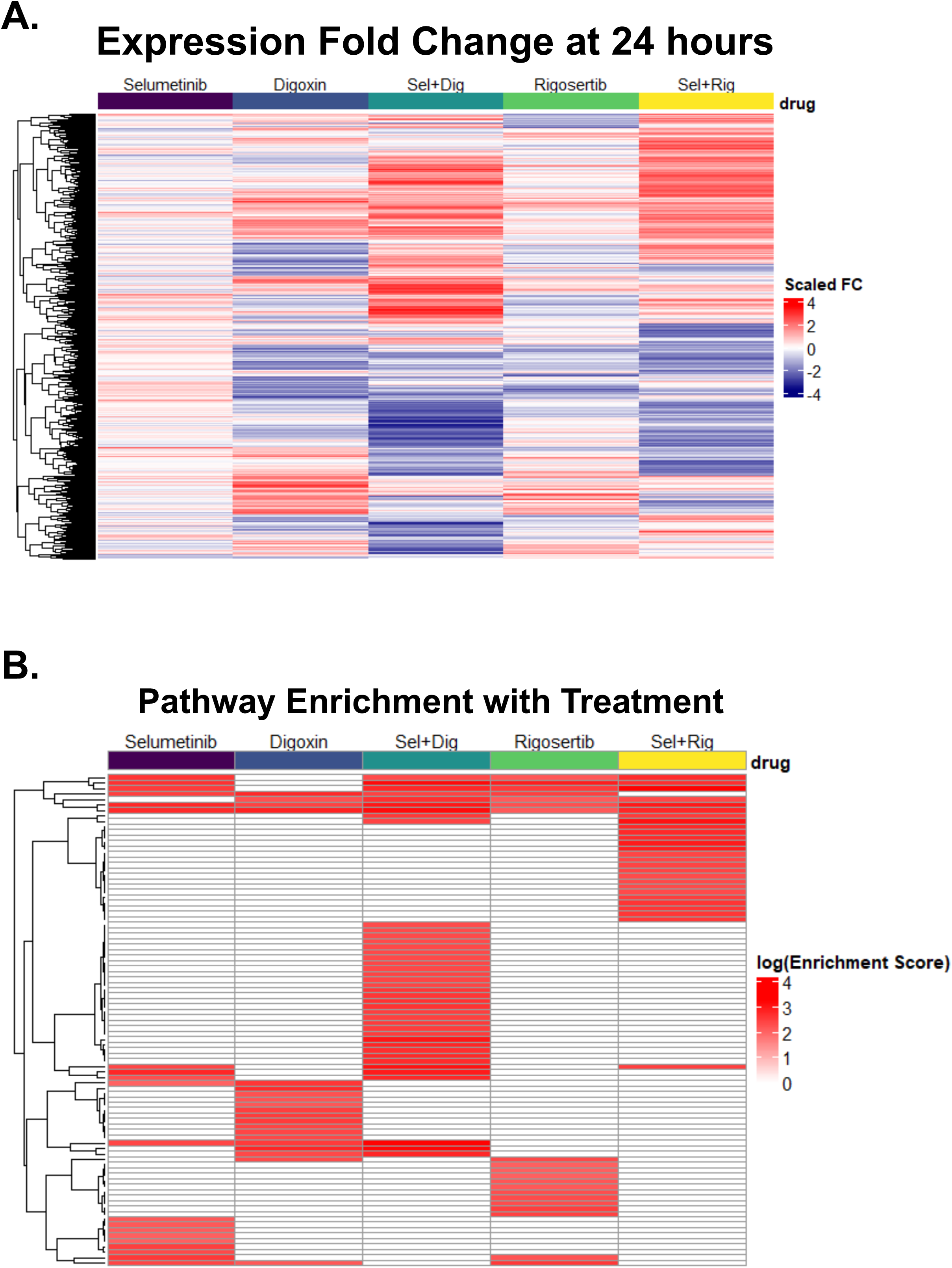
Cellular responses to single agent and combination drug therapies. A. mRNA expression profile of S462TY cells treated for 24hrs with digoxin, rigosertib, selumetinib, digoxin + selumitinib and rigosertib + selumetinib. B. Combination drug treatment results in unique emergent cellular responses not seen in single agent therapies. Pathway enrichment as determined by SuperPath analysis on RNAseq data obtained from S462-TY cells subjected to drug treatment for 24hrs (single and combo).

Gene set enrichment analysis was performed on the transcriptome data to distill the biological responses observed in each drug treatment condition, identifying shared and uniquely effected pathways (Figure 5B). Column 3 of Figure 5B corresponds to the selumetinib + digoxin combination treatment. A large cluster of unique enriched pathways can be seen in the center of the plot. These pathways include GPCRs, Akt signaling, further enrichment of Erk signaling components, members of the cardiac conduction pathway (likely driven by an increased calcium channel perturbation in the combination treatment), the CREB pathway, Interleukin-10 signaling, and others (Supplementary Figure 3C and 5). Interestingly, selumetinib treatment alone resulted in dysregulation of ion channel transporters, including calcium and potassium transporters (Supplementary Figure 3A). Digoxin inhibits the Na+/K+-ATPase pump, thus leading to an increase in intracellular calcium, and digoxin treated cells upregulated potassium channel transcripts, likely as a compensatory mechanism in response to this insult (Supplementary Figure 3B). The ion channel homeostasis dysregulation driven by Selumetinib, combined with the direct inhibition of the Na+/K+-ATPase pump by digoxin, could partially explain the synergy seen with this drug combination against MPNST.

The combination treatment of selumetinib + rigosertib also elicited novel enriched pathways not observed in the corresponding single drug treatments (Figure 5B, column 5, Supplementary Figure 4B). Enriched, regulated, pathways of note include several immune system response components such as interleukin 9 signaling, cytotoxic T-cell mediated apoptosis, antigen presentation, CD28 co-stimulation, and other immune response signatures (Supplementary Figure 4C and 5). These responses suggest the selumetinib + rigosertib combination might perform even better in an immune competent *in vivo* model or in the clinic, where the host immune system could be engaged.

### In vivo activity of small molecules identified in the high-throughput drug screen

We tested the efficacy of 4 of the top hits from our high-throughput drug screening effort, and the MEK inhibitor selumetinib, in a genetically engineered mouse model (GEMM) of high-grade peripheral nerve sheath tumors. Using a Cre recombinase driven by the Desert hedgehog promoter with *Nf1^fl/fl^, Pten^fl/fl^*, this model develops multifocal disease in all major peripheral nerves, including dorsal root ganglia, brachial plexus, sciatic, and trigeminal nerves, with 100% penetrance^32^. Untreated animals develop severe ataxia and become moribund prior to weaning, at roughly 18 days of age.

For *in vivo* testing to have increased translation relevance, two of the small molecules identified in the initial screening work were substituted for derivatives that have undergone further clinical development to increase their drug-like properties. The PP2A inhibitor LB100 is a cantharidin derivative currently undergoing multiple clinical trials, and dimethylamino parthenolide (DMAPT) is an orally bioavailable parthenolide analogue^51,52^. All of the single agent drugs significantly extended the lifespan of the animals in DhhCre;*Nf1^fl/fl^, Pten^fl/fl^* mice (Figure 6A, Table 4). Vehicle treated animals had a median survival of 18 days, whereas the cohorts treated with rigosertib (22 days), LB100 (24 days), digoxin (27 days), DMAPT (27.5 days), and selumetinib (24 days) showed increased survival.

**Figure 6.**
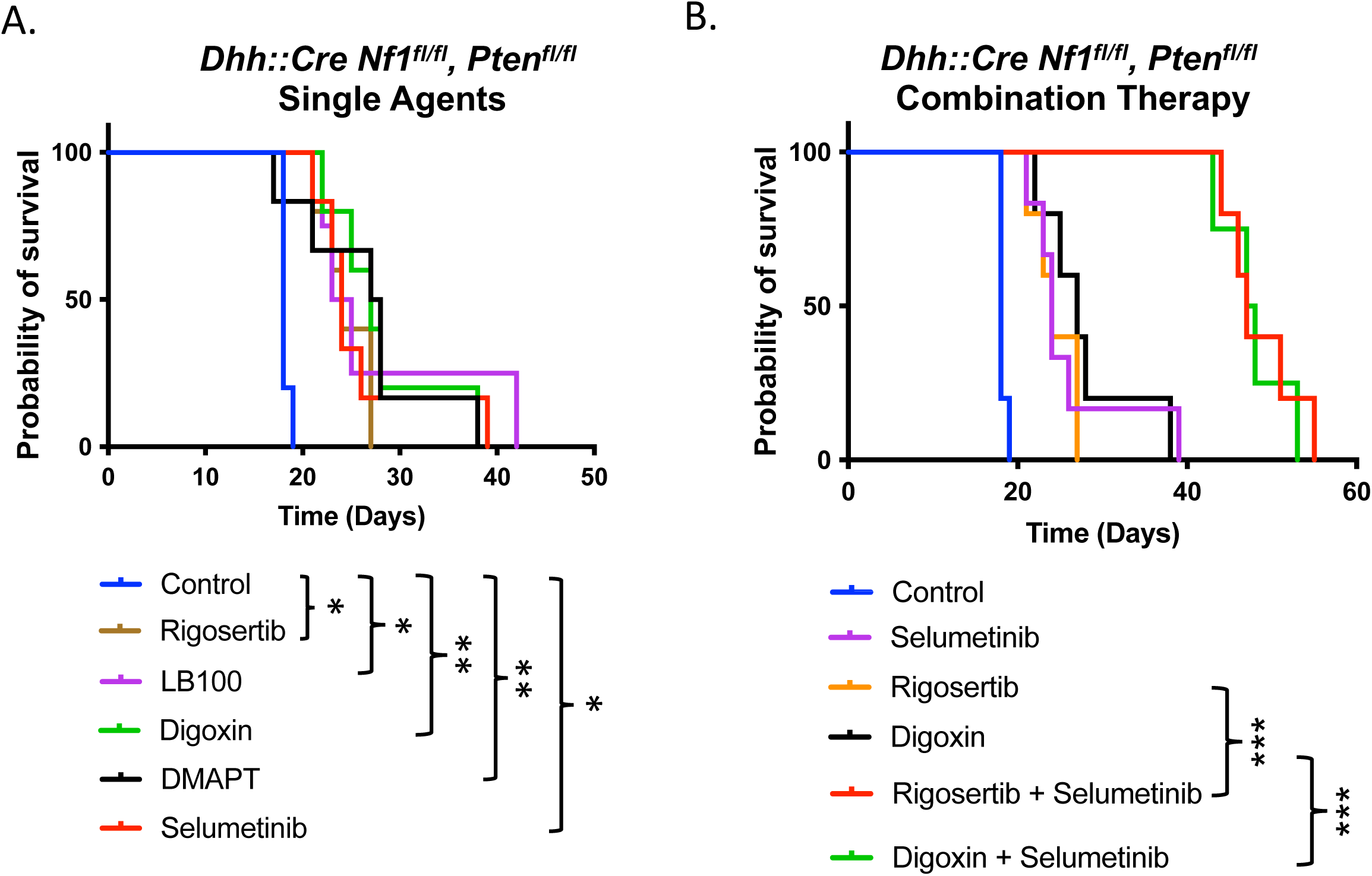
Drugs identified in our HTS are effective at prolonging life in a rapidly progressing genetically engineered mouse model (GEMM) of high-grade peripheral nerve sheath tumors. A. *Dhh::Cre, Nf1^fl/fl^, Pten^fl/fl^* mice were randomized to control or treatment at 7 days of age. All single agent therapies increased survival in this model. B. Combining digoxin or rigosertib with the MEK inhibitor selumetinib significantly increased overall survival compared to both control and monotherapies. * *p*<0.05 ** *p*<0.01, *** *p*<0.001.

**Table 4.**
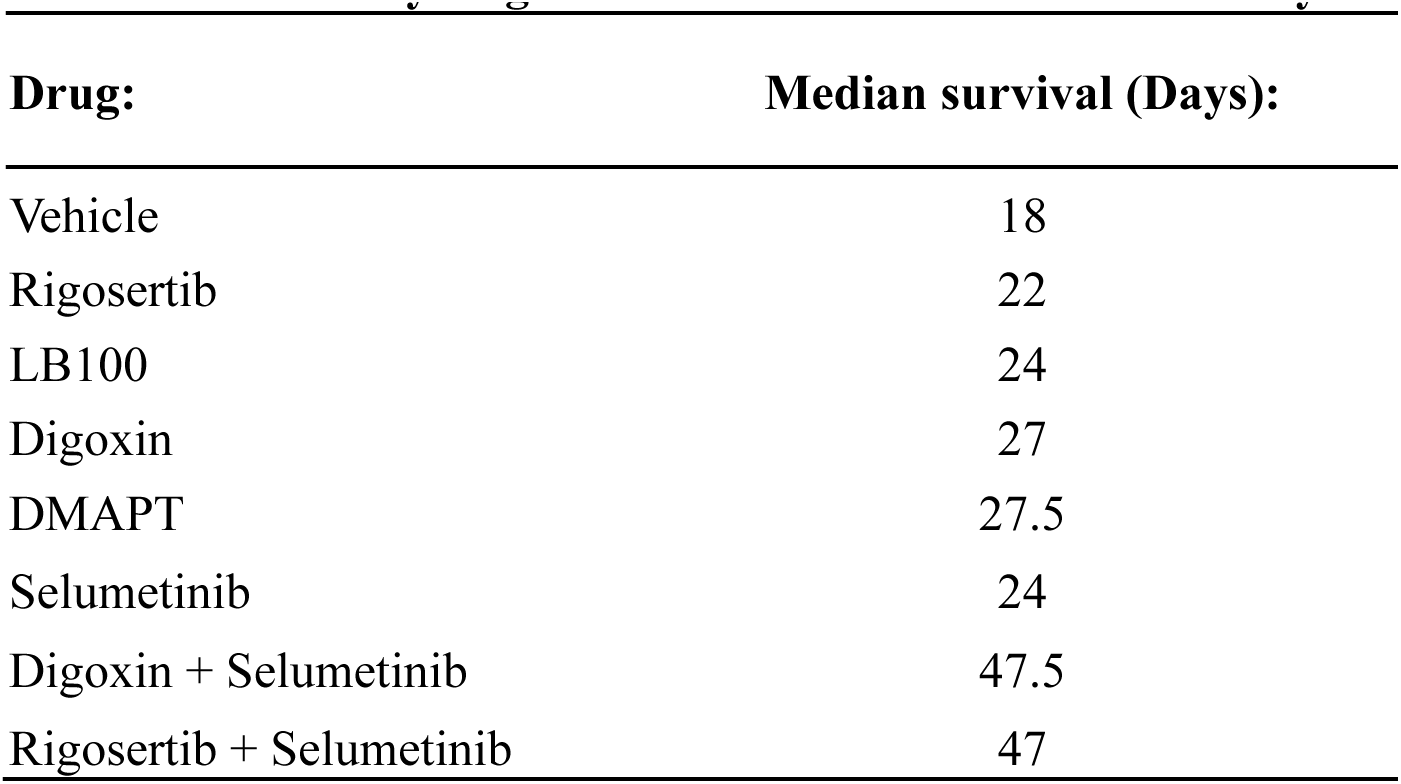
Genetically Engineered Mouse Model Survival Analysis.

| Drug: | Median survival (Days): |
| --- | --- |
| Vehicle | 18 |
| Rigosertib | 22 |
| LB100 | 24 |
| Digoxin | 27 |
| DMAPT | 27.5 |
| Selumetinib | 24 |
| Digoxin + Selumetinib | 47.5 |
| Rigosertib + Selumetinib | 47 |

Given that digoxin is FDA approved for other indications and rigosertib is currently in several late-stage clinical trials, and due to the strong *in vitro* synergy of their combination effects in MPNST cell lines, these agents were deemed of particular interest. Therefore, combination therapy in conjunction with the MEK inhibitor Selumetinib was used in the DhhCre;*Nf1^fl/fl^, Pten^fl/fl^* mice GEMM. The rigosertib + selumetinib and digoxin + selumetinib combinations each showed dramatic increases in median survival, when compared to their single agent results (22 day to 47 days and 27 days to 47.5 days, respectively, *p*<0.001) (Figure 6B, Table 4). These increases in survival are among the largest observed in this *in vivo* model.

To determine the suitability of the drugs identified in this screening effort for the potential use against MPNST *in vivo*, the same five compounds used in the GEMM were tested against a human MPNST cell line xenograft (S462-TY) grown as flank tumors in immunodeficient NRG mice. Tumors in vehicle treated mice progressed rapidly, after reaching the study size enrollment of ∼200mm^3^, with tumors reaching end point size criteria of 2000mm^3^ in about a week. All the single agents improved survival significantly in this model (Figure 7A). However, DMAPT significantly underperformed compared to the other single agents when tested against this MPNST xenograft. While all the single agents tested slowed tumor growth, some of the drugs were able to shrink established tumors, with digoxin and LB100 having the strongest tumor lytic activity (Figure 7B). In fact, digoxin was able to rapidly shrink some tumors to a nearly undetectable size. However, the xenograft tumors eventually escaped each of the tested monotherapies and progressed to endpoint, necessitating euthanasia.

**Figure 7.**
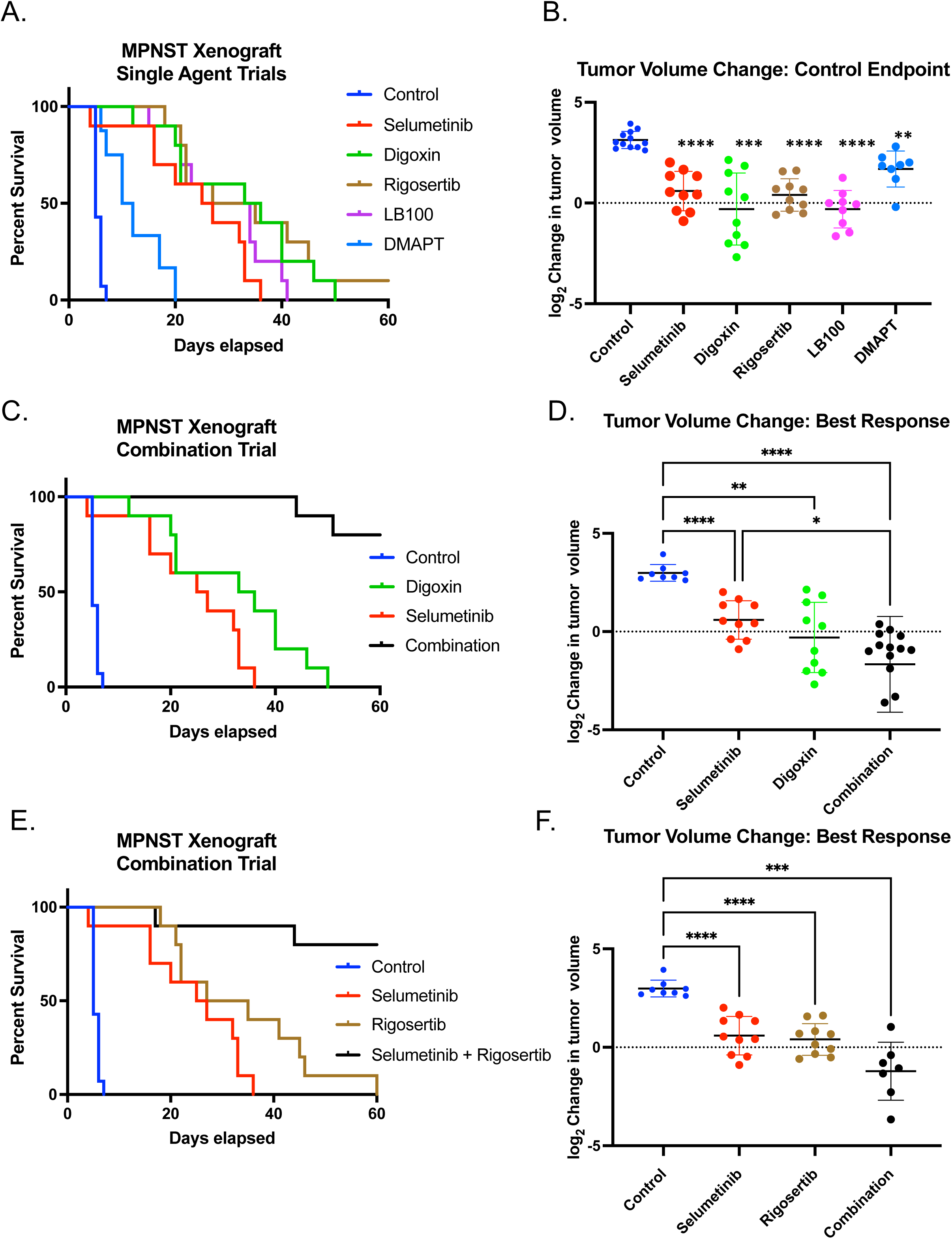
Drugs identified in our HTS are effective against human MPNST xenograft models, increasing survival and causing tumor regression. S462-TY MPNST xenografts were established in NRG immunodeficient mice. Animals were randomized into control and treatment groups when tumors reached ∼200mm^3^. Animals were euthanized when tumors reached 2000mm^3^. A. Single agent therapy controlled tumor growth and prolonged overall survival. B. Log_2_ fold change of tumor volume in control and treated tumors. All monotherapies controlled tumor growth, with digoxin reliably able to significantly shrink tumors. ** *p*=0.0097, *** *p*=0.0007, **** *p=<0.0001* C. Digoxin and selumetinib combination treatment dramatically increased survival compared to single agent therapy, with long term survivors at the trial end. D. Nearly all tumors treated with digoxin/selumetinib combination showed tumor regression, with some having a complete response. * *p*=0.0326, ** *p*=0.0013, **** *p=<0.0001* E. Rigosertib and selumetinib combination treatment dramatically increased survival compared to single agent therapy, with long term survivors at the trial end. F. Tumors treated with rigosertib/selumetinib combination showed tumor regression, while single agent treatment largely results in stable disease. *** *p*=0.0005, **** *p=<0.0001*

The combination of digoxin + selumetinib, which showed synergy when tested in the GEMM, was therefore tested in the S462-TY MPNST cell line xenograft. A dramatic effect was seen when combination therapy was tested in this model. Not only did the combination significantly outperform either of the single agents, but we also observed long term survivors at the conclusion of the study (Figure 7C). Some animals had no detectable tumors at the conclusion of the study, suggesting a durable response. Moreover, while digoxin alone shrank some of the tumors, the majority of the tumors treated with the digoxin + selumetinib combination showed rapid shrinkage with marked reduction in tumor size (Figure 7D). Tumors from mice treated with this combination also showed extensive TUNEL positivity and cleaved Caspase 3 via IHC (Supplementary Figure 6). We also tested the selumetinib + digoxin combination in an MPNST patient-derived xenograft (PDX) grown in NRG mice. In the PDX, an increase in survival and reduction in tumor growth was also observed (Supplementary Figure 7), although it was not as robust as the response seen in the GEMM or MPNST cell line xenograft.

Since the combination of rigosertib + selumetinib showed promise as a synergistic combination in the *in vitro* models tested in this study and the GEMM model of high grade PNST described earlier, it was also tested as combination therapy in the MPNST cell line xenograft. Much like the combination of digoxin + selumetinib, a dramatic improvement in survival was seen in tumor bearing animals treated with this combination (Figure 7E). The response was improved over each matched monotherapy, and long term survivors were observed at the end of the study. This combination was also not simply tumor static; all but one tumor bearing animal showed tumor shrinkage during the therapeutic trial (Figure 7F). Likewise, when the selumetinib + rigosertib combination was tested in the MPNST PDX mentioned, control of tumor growth was superior to either monotherapy, and there was a corresponding increase in overall survival (Supplementary Figure 8). The response of this combination in the PDX model was significantly better than the selumetinib + digoxin combination.

## Discussion

The management of Neurofibromatosis Type 1 (NF1)-associated Malignant Peripheral Nerve Sheath Tumors (MPNSTs) remains a difficult clinical challenge, necessitating innovative therapeutic strategies. Recent breakthroughs in cancer research underscore the potential of synthetic drug sensitivity and combination therapy approaches in addressing the intricacies of NF1-associated tumors, particularly the dysregulation of the Ras-MAPK pathway. The identification and exploitation of synthetic lethality relationships within these tumors could pave the way for precision medicine tailored to the specific molecular vulnerabilities of NF1-associated MPNSTs^13,53^.

MEK inhibitors, such as selumetinib, have demonstrated efficacy in targeting the aberrant Ras-MAPK signaling pathway observed in NF1-associated plexiform neurofibromas, but have not been useful for treating most MPNSTs^29,54^. Synthetic drug sensitivity, grounded in the concept of exploiting cancer cell-specific vulnerabilities, could become a critical element in optimizing therapeutic interventions like MEK inhibition^55^. However, tumors can evolve avenues to bypass MEK dependency, such as upregulation of receptor tyrosine kinases^56^. So, while important, MEK inhibition is not sufficient on its own for MPNST treatment, motivating our search for new drugs useful in *NF1*-deficient tumors.

Here we report several novel drugs, and mechanisms of action, that are selectively lethal to *NF1*−/− human Schwann cells and MPNST. Some classic chemotherapeutic agents did show selective killing against *NF1-*deficient cells, particularly topoisomerase inhibitors which have been used in the clinic for management of MPNST^57^. While we did identify some small molecules with targets in the Ras-MAPK pathway, most of the hits were for targets outside of this signaling axis that would have been difficult to predict before screening. These include modulators of the NF-κB pathway, agents that contribute to ER stress and the UPR, and a large set of compounds that increase the intracellular levels of calcium. The unexpected classes of selective agents are particularly interesting since most have not been pursued for clinical translation for *NF1*-deficient tumors and can provide insight into their underlying biology. Thus, our screening methodology shows the strength of using a diverse collection of small molecules to uncover druggable *NF1-* related synthetic lethal interactions.

The class of molecules identified which impose ER stress upon treatment are of note (Table 2). Cantharidin is a blister agent and poison at high doses, but a derivative has been developed clinically as an inhibitor of the serine/threonine phosphatase, protein phosphatase 2A (PP2A)^58^. PP2A has was originally considered a tumor suppressor, because of its role counteracting kinase-driven signaling pathways and maintaining intracellular homeostasis and the fact it is found mutated in cancers driven by oncogenic kinases, such as chronic myelogenous leukemia^59^. However, inhibition of the phosphatase with the small molecule LB100 has been found effective in numerous preclinical models of a diverse set of cancers, including nervous system tumors and sarcomas^52,58,60^. Cantharidin was also identified in a previously reported medium-throughput drug screening project to identify compounds selective against *Nf1-*deficient mouse embryonic fibroblasts^20^. However, the effect of the compound in that system was modest *in vitro* and it was not tested by *in vivo* validation. This previous report corroborates our findings, and we see a moderate *in vivo* response to PP2A inhibition in both a GEMM of high grade PNST and in MPNST cell line xenografts. LB100 has been clinically developed and is in Phase I and II clinical trials for both solid tumors and myelodysplastic syndrome^61^ (NCT03886662). This class of compound is likely providing a therapeutic benefit in NF1-associated models via two mechanisms. First, there is already heightened Ras-MAPK and Ras-PI3K signaling flux in these peripheral nerve sheath tumors. Inhibiting an important phosphatase responsible for keeping the pathways in check could lead to their hyperactivation and induction of apoptosis, or cell cycle arrest as has been described in other contexts^62^. PP2A inhibition with LB100 is also a strong chemo and radiosensitizer in many cancer models^58^. Secondly, cantharidin and LB100 are known to cause ER stress and induce the unfolded protein response in cells. The UPR and ER stress is already activated in neurofibromagenesis through Runx1/3-driven adaptive response^63^. Given that the UPR can easily tip from supporting cell survival to inducing apoptosis, additional ER and UPR stress in these cells, induced by LB100 treatment, could drive this pathway toward cell death.

Several small molecules influencing the NF-κB pathway were also identified in this screening effort, with one subsequently followed up on with *in vivo* testing. Parthenolide is a sesquiterpene lactone natural product from plants, with high anti-inflammatory and antitumor bioactivity^64,65^. While parthenolide has poor bioavailability, several analogues have been developed to be orally bioavailable and show promise as *bona fide* drugs for cancer treatment^51,66^. This class of compounds has been shown to target the NF-κB pathway, resulting in downregulation of transcriptional targets^67,68^. This pathway is emerging as important for neurofibroma development, with a report showing NF-κB dysregulation in multiple *Nf1−/−* Schwann cell populations in a genetically engineered mouse model of plexiform neurofibroma development^69^. Parthenolide also induces reactive oxygen species (ROS)-mediated apoptosis in cancer models^70^. Other drugs identified in this study, including the calcium level modulators, can increase cellular ROS levels. Catastrophic oxidative stress has been demonstrated as a vulnerability in RAS-driven cancers, including MPNST^71^, supporting the idea that compounds we have identified as converging on this cellular stress axis have a biological rationale.

Most of the small molecules identified in this study target pathways outside of the canonical MAPK signaling axis, and some show marked evidence of synergy when combined with MEK inhibition. Two novel drug combinations exhibit high synergism with the MEK inhibitor selumetinib, significantly enhancing its therapeutic efficacy in multiple models of MPNSTs. Given that MEK inhibition is widely used for the treatment of symptomatic plexiform neurofibromas and that most MPNSTs arise from preexisting plexiform neurofibromas, compounds identified to work well when combined with MEK inhibition are of high translational interest^13,30,72^.

The first synergistic combination we explored comprises the MEK inhibitor selumetinib and digoxin, a cardiac glycoside with emerging anti-cancer properties. The rationale for this combination is grounded in the potential of digoxin to modulate cellular signaling, particularly by targeting the Na+/K+-ATPase pump. Digoxin may disrupt the cellular microenvironment, rendering MPNST cells more susceptible to the inhibitory effects of selumetinib^73–75^. The increased intracellular calcium resulting from digoxin treatment can add mitochondrial and oxidative stress to the cells, to both of which *NF1*-deficient cells show an increased susceptibility^71^. Furthermore, pathway analysis of RNAseq data in this study shows that human MPNST cells treated with the MEK inhibitor selumetinib have altered expression of ion channel genes (Supplementary Figure 3A). This suggests dysregulated ion homeostasis is already occurring upon MEK inhibition. The addition of further ionic stress by digoxin to this already dysregulated state could push the cells over a tipping point, resulting in the strong therapeutic response we see both *in vitro* and *in vivo* with combination selumetinib/digoxin treatment.

There are potential safety concerns when using a cardiac glycoside, such as digoxin, for cancer therapy. However, this class of drugs has been used for decades and the clinical management is very well described in both the pediatric and adult populations^76,77^. There is also a fast-acting antibody-based antidote to reverse acute effects of digoxin^78^. The combination of digoxin plus the MEK inhibitor trametinib has been shown to achieve disease control in metastatic melanoma patients in a phase 2 clinical trial^74^. The synergy seen in our current study, combined with success seen using this combination in other cancer types, suggests combination treatment with a MEK inhibitor and digoxin to be a viable option for the treatment of NF1-associated peripheral nerve sheath tumors.

The second combination involves the MEK inhibitor selumetinib paired with rigosertib, which was developed as a dual PLK1 and PI3K inhibitor^79^. The rationale behind this combination lies in the complementary inhibition of multiple signaling pathways implicated in MPNST progression. The combined inhibition of both the Ras-MAPK and PI3K pathways may lead to a more comprehensive blockade of pro-survival signals, potentially overcoming compensatory mechanisms that limit the effectiveness of single-agent therapies^80^. There is some uncertainty regarding the true cellular target of rigosertib at achievable human doses, with strong evidence suggesting it is a microtubule destabilizing agent^31^. One well designed study reports rigosertib’s ability to induce mitotic arrest in RAS-mutated sarcomas and neuroblastoma, though its impact on mitotic the spindle. In that study, rigosertib also behaved synergistically when used in combination with the MEK inhibitor trametinib^81^, as in our models of NF1-associated tumors, which also have hyperactive Ras signaling. Using pathway enrichment analysis on MPNST cells treated with the MEK inhibitor selumetinib and rigosertib we determined there are a marked number of immune responsive components modulated in response this this combination, including antigen presentation. This is intriguing as it could provide a way to engage the host immune system against MPNSTs, in addition to the direct tumor cell killing effect of the combination. To date rigosertib has undergone extensive clinical testing, including Phase I and Phase III trials, for a variety of malignancies where it has shown efficacy^47,82^. The strong activity and synergy seen in our models would suggest this is a prime candidate for the treatment of NF1-associated tumors, particularly MPNST, when combined with MEK inhibition.

The establishment of validated cell-type specific screening platform that can identify small molecule hits which show effectiveness across several models of NF1-associated neoplasia is an exciting and adaptable development for the field. While this current iteration of the system introduces loss of function mutations in *NF1* into human Schwann cells that were immortalized with human *TERT* and expression of an activated murine *Cdk4* (thus mimicking an atypical-like neurofibroma genotype), known mutations associated with progression to MPNST could be engineered into these cell-line models and further emergent synthetic drug sensitivities identified. Candidates for this next step would be introducing loss of function mutations in the polycomb repressor complex 2 (PRC2) components, such as *EED* or *SUZ12*, since ∼80% of MPNST are PRC2-deficient^83,84^. The same platform can also be adapted to conduct synthetic lethal genetic screens for *NF1* synthetic lethal partners, using genome wide CRISPR knockout libraries^13,26,85^. Studies such as these would provide a further unbiased approach to discover novel drug targets and new biology about the function of *NF1* in Schwann cells.

The identification of novel drug combinations, selumetinib with Rigosertib and Selumetinib with digoxin, represents a promising avenue for the treatment of NF1-associated peripheral nerve sheath tumors including PNs, ANFs and MPNSTs. These combinations offer a synergistic approach by concurrently targeting distinct signaling pathways, thereby potentially enhancing the therapeutic efficacy and overcoming limitations associated with single-agent treatments. Further preclinical and clinical investigations are warranted to validate the safety and efficacy of these combinations, with the goal of improving outcomes for patients with this aggressive and challenging malignancy.

## Acknowledgements

The authors would like to thank Dr. Margaret Wallace for providing iHSCs and Dr. Nancy Ratner for providing MPNST cells lines. We acknowledge Dr. James Walker for important advice and discussions during planning of this work.

## Authorship statement

Designed the study: K.B.W., D.A.L. Advised the study: A.T.L., B.J.K., T.J., N.R., C.L.M., D.A.L. Acquired funding: D.A.L, N.R., K.B.W. Acquired and generated cell lines: K.B.W, A.T.L., R.L.W. Performed experiments and analyzed data: K.B.W., B.J.K., A.T.L., K.E.C., R.L.W., W.A.H. Performed bioinformatic and statistical analysis: T.J. Wrote the manuscript: K.B.W.

