## Supplementary Material for "Pharmacogenomic synthetic lethal screens reveal hidden vulnerabilities and new therapeutic approaches for treatment of NF1-associated tumors"

### Supplementary Information

**Measurement of RAS-GTP levels.** Cells were harvested at 80% confluency, counted using the Countess II (Life Technologies), and plated with DMEM with 1% pen/strep and 10% FBS. The following day, media was replaced with DMEM with 1% pen/strep and no FBS, and cells were serum starved overnight. At that time cells were harvested, washed with cold PBS and pellets kept on ice prior to processing. The Active Ras Pull-Down and Detection Kit (Thermo Scientific) was then used according to the manufacturer's protocol. Briefly, cells were lysed in 500μL Lysis/Binding/Wash Buffer containing complete protease inhibitor cocktail (Roche). RAS-GTP was extracted by lysates by affinity precipitation with GST-Raf1-RBD, and eluted in 50μL 2x reducing sample buffer (1 part β-mercaptoethanol to 20 parts Sodium dodecyl sulfate (SDS) Sample Buffer). Samples were heated for 5 min at 95°C, and 25μL used for western blot analysis.

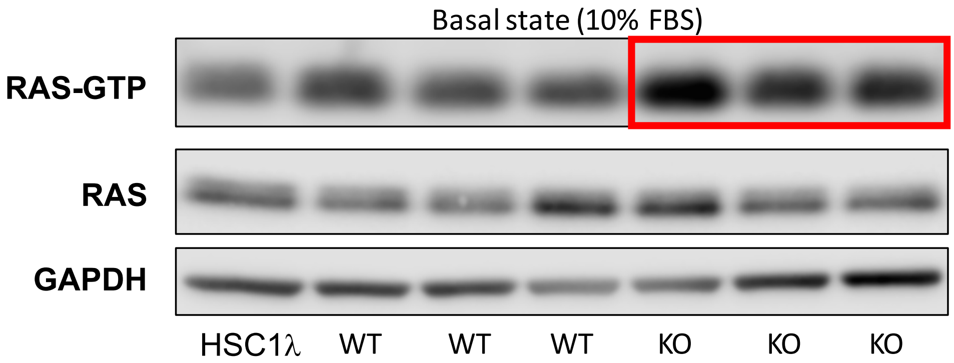


**Supplementary Figure 1.** *NF1*-deficient iHSCs display elevated basal levels of active RAS (RAS-GTP) compared to *NF1*-proficient isogenic matched sister clones and the parental cell line, HSC1λ. Lysates were processed using an active RAS pull-down kit and probed via immunoblot with an anti-RAS antibody in the RAS-GTP fraction (top) or total RAS within the cell lysate (middle). GAPDH was probed as a cell loading control (bottom).

***In vivo* tumor forming ability of iHSCs in NRG mice.** *NF1*-proficient and *NF1*-deficient engineered clones were injected into the flanks of 6 to 7-week old NOD-Rag1null IL2rgnull, NOD rag gamma, NOD-RG (NRG) mice ($1.0\times10^6$ cells) in 50% Matrigel (BD Biosciences). Mice were monitored for tumor formation for 6 months post injection or until a tumor volume of 2000mm$^3$ was reached. The human MPNST cell line S462-TY was used as a positive control for tumor formation. Tumors were defined as a mass reaching a volume of at least 150mm$^3$. Summary shown in Supplementary Table 1.

**Supplementary Table 1.**
***In vivo* growth of immortalized human Schwann cells and MPNST in NRG mice.**

| Injected Cell Line | Genotype | Mice Injected | Tumors Formed |
|---|---|---|---|
| HSC1λ | Immortalized human Schwann cell line NF1 +/+ | 5 | 1 |
| S462TY | Human MPNST cell line | 4 | 4 |
| N1 (3) | NF1 +/+ | 8 | 0 |
| N1(10) | NF1 +/+ | 4 | 0 |
| N0 (32) | NF1 -/- | 4 | 4 |
| N0 (5) | NF1 -/- | 5 | 5 |

Tumors were defined as a tumor volume of at least 150 mm$^3$. Mice were monitored six months post injection or until tumor volume reached 2000 mm$^3$.

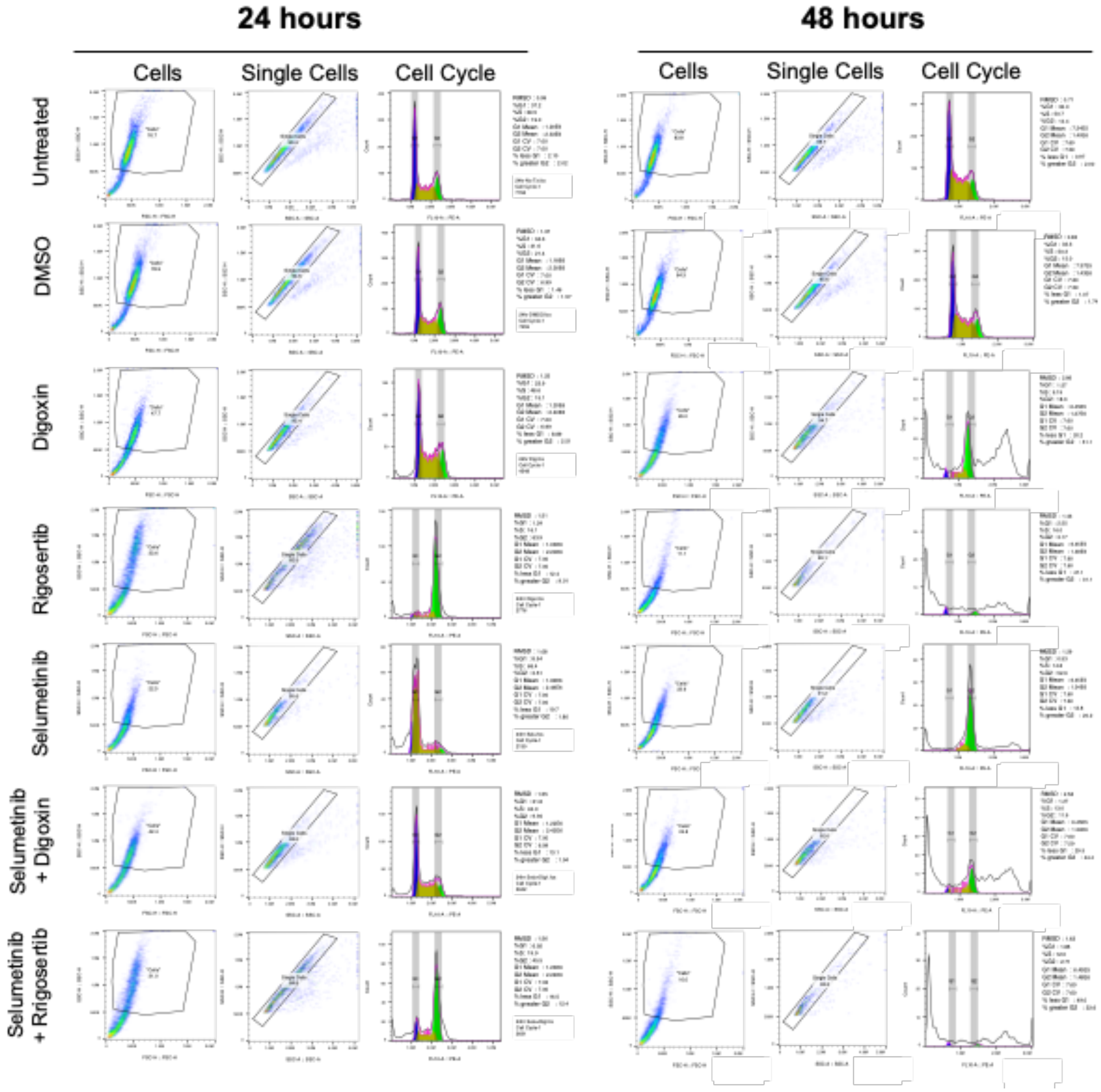


**Supplementary Figure 2.** Cell cycle analysis of S462-TY MPNST cell line in response to drug treatments. Cells were exposed to drug treatment, vehicle, or untreated (as indicated) for 24 or 48 hours. Cell cycle analysis was completed using a Propidium Iodide kit (Abcam ab139418) using manufacturer's protocol as described in Materials and Methods. Gating strategy for identify cell population fraction, single cells, and finally cell cycle determination are indicated above, for each condition.

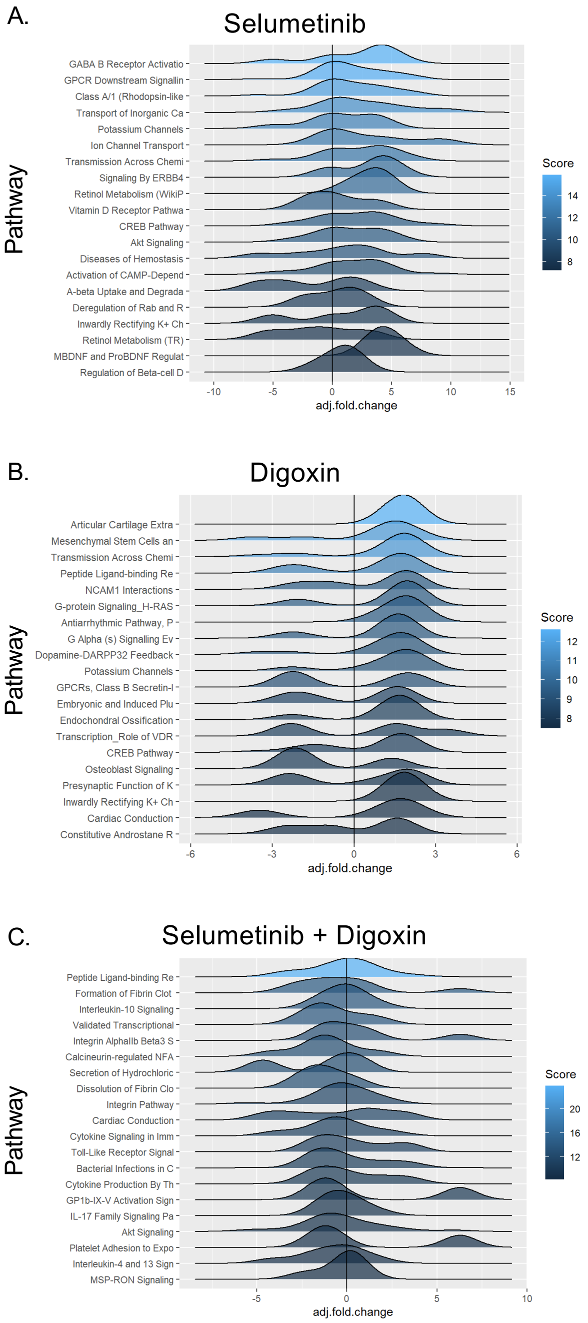


**Supplemental Figure 3.** Perturbation of enriched pathways upon drug treatment of S462Ty cells with: A. Selumetinib, B. Digoxin, C. Selumetinib + Digoxin. Cells were exposed to vehicle (DMSO) or the indicated drug for 24 hrs, at which time mRNA was harvested and subjected to Next-generation RNA sequencing. Pathways determined by SuperPath analysis pipeline. Color indicates enrichment score. Height of peak represents number of pathway members enriched. Adjusted fold change is relative fold change of treatment condition compared to vehicle exposure.

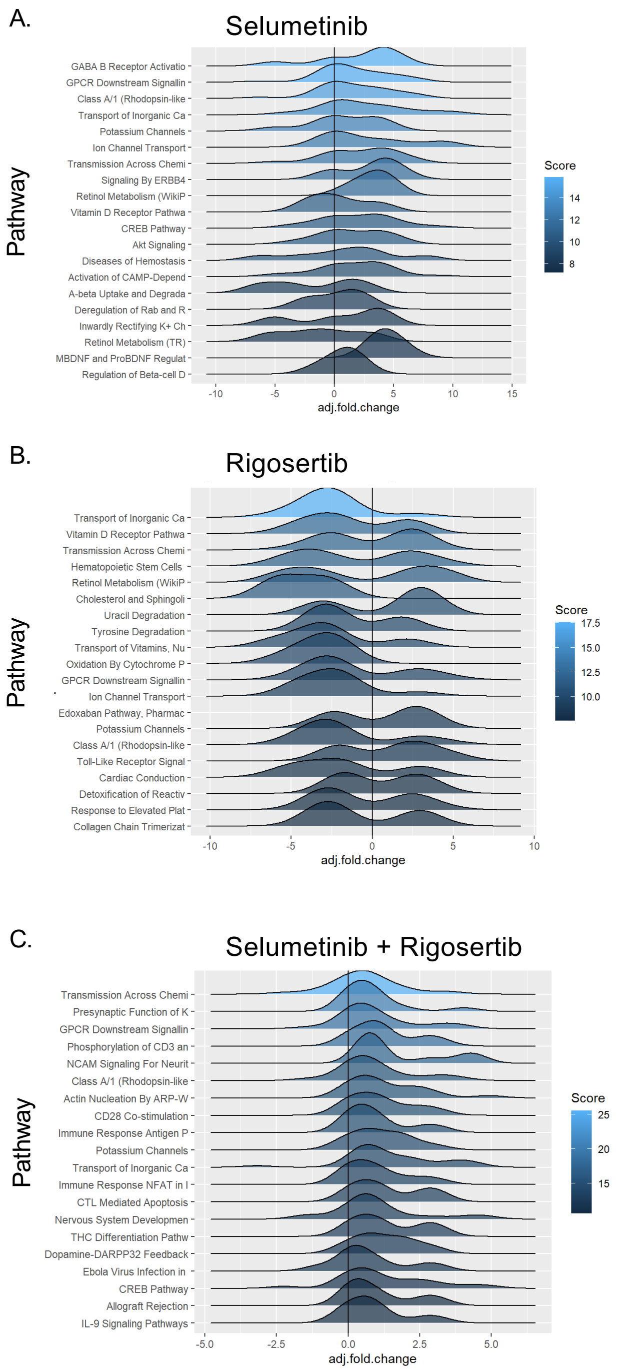


**Supplemental Figure 4.** Perturbation of enriched pathways upon drug treatment of S462TY cells with: A. Selumetinib, B. Rigosertib, C. Selumetinib + Rigosertib. Cells were exposed to vehicle (DMSO) or the indicated drug for 24 hrs, at which time mRNA was harvested and subjected to Next-generation RNA sequencing. Pathways determined by SuperPath analysis pipeline. Color indicates enrichment score. Height of peak represents number of pathway members enriched. Adjusted fold change is relative fold change of treatment condition compared to vehicle exposure.

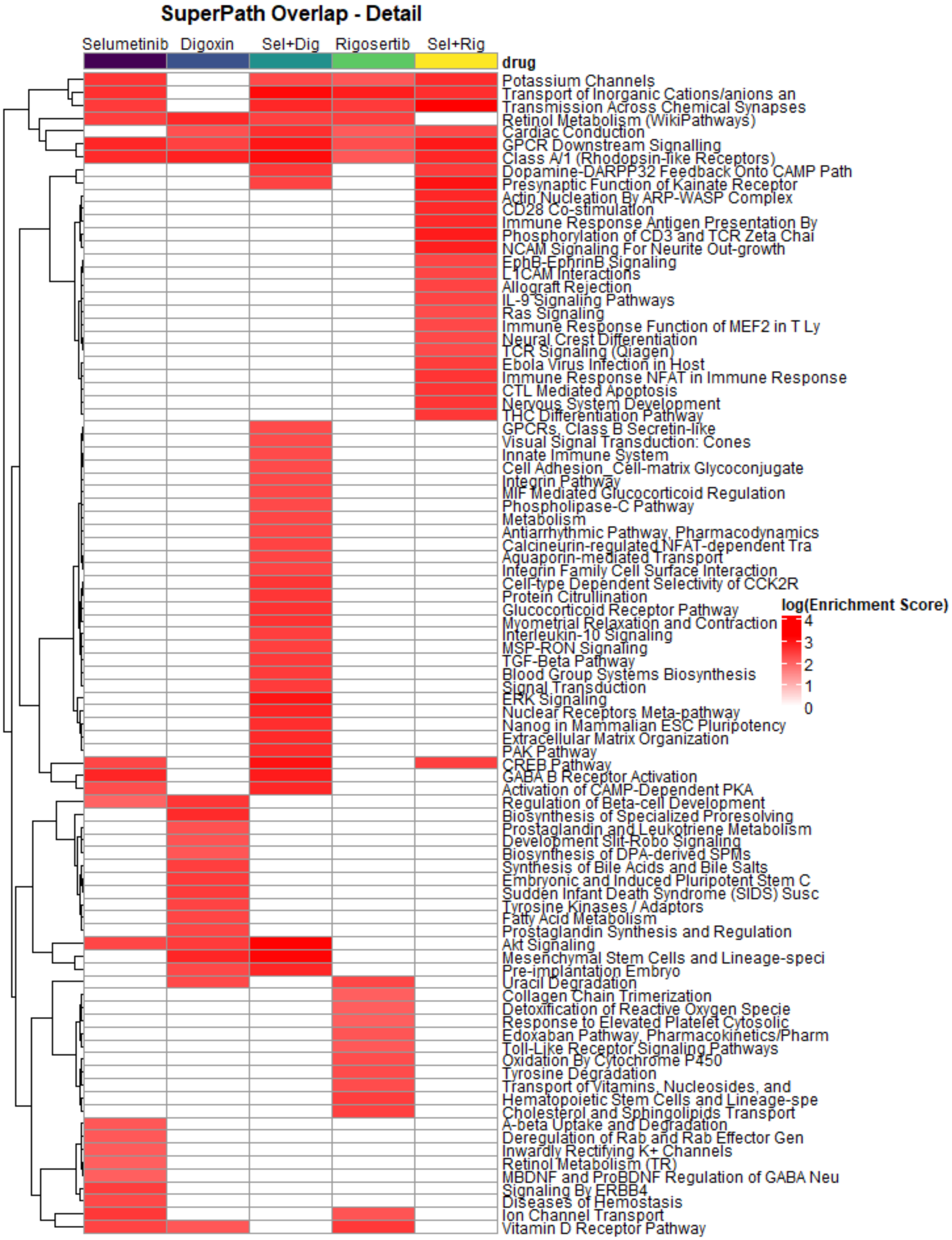


**Supplementary Figure 5.** Pathway enrichment analysis of drug treated cell, as determined by SuperPath. This is a duplicated of Figure 5 of the main text, with the individual pathways listed.

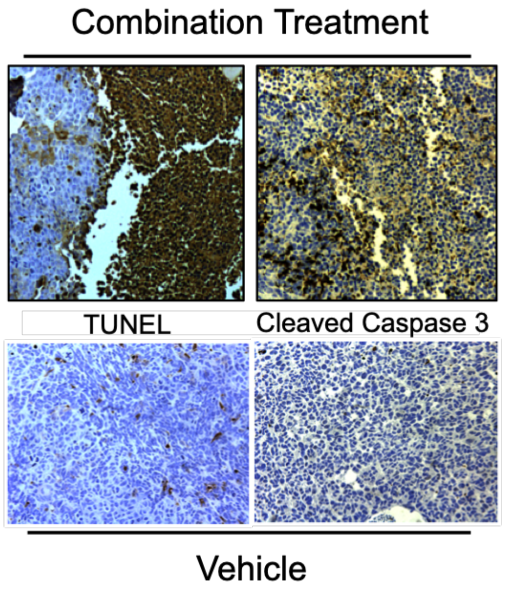


**Supplementary Figure 6.** The combination of digoxin + selumetinib causes cell death and induction of apoptosis in MPNST xenograft tumors. Human MPNST cell line S462-TY xenograft was established in the flanks of NRG mice and allowed to grow to a size of ~500mm$^3$. Tumor bearing animals were then treated with the drug combination or vehicle for 5 days. Tumors were harvested, processed into formalin fixed, paraffin embedded blocks and stained for TUNEL positivity and cleaved caspase 3.

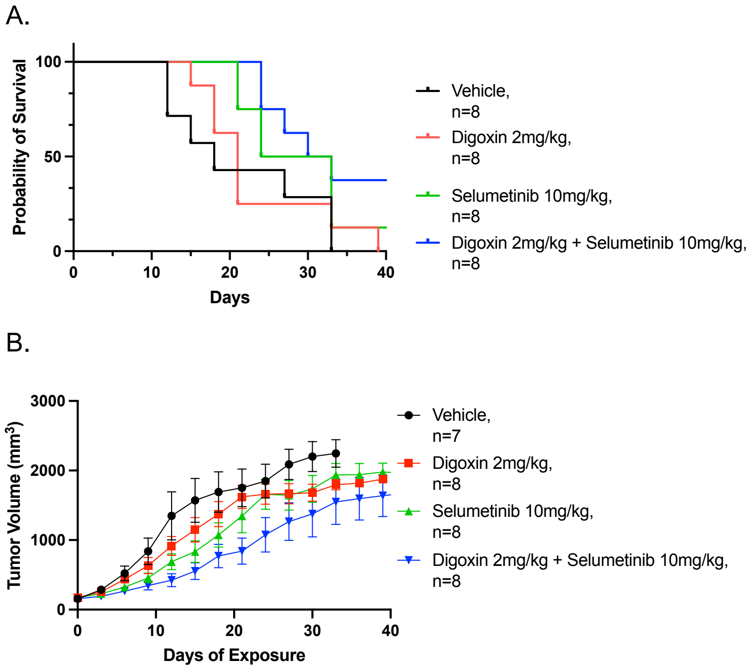


**Supplementary Figure 7.** Response of an MPNST PDX to treatment with selumetinib, digoxin, or the combination. A. Kaplan Meier survival analysis of each cohort corresponding to these treatments. B. Tumor growth kinetics with vehicle or the indicated treatment.

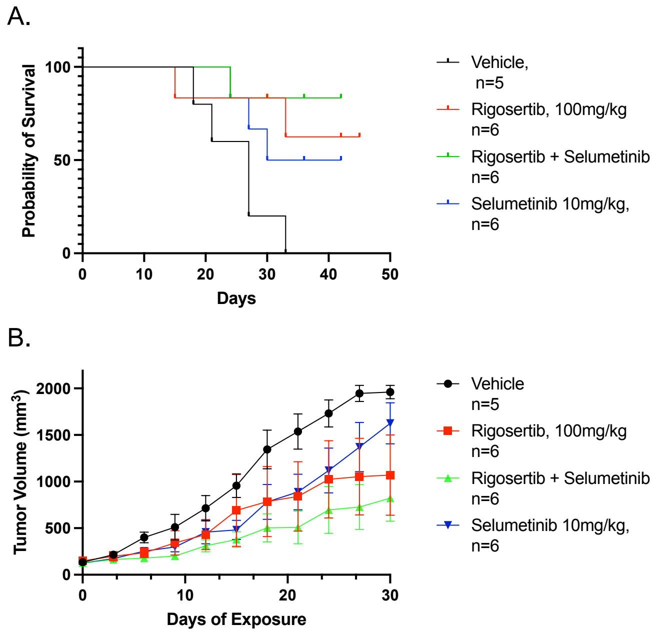


**Supplementary Figure 8.** Response of an MPNST PDX to treatment with selumetinib, rigosertib, or the combination. A. Kaplan Meier survival analysis of each cohort corresponding to these treatments. B. Tumor growth kinetics with vehicle or the indicated treatment. Combination treatment with rigosertib and selumetinib provided a substantial improvement in tumor control and overall survival in this PDX model.
